# Temporal single-cell analysis reveals age-associated delay in immune resolution after respiratory viral infection

**DOI:** 10.64898/2025.12.15.694321

**Authors:** Yue Wu, Chaofan Li, Jinyi Tang, Xiaochen Gao, In Su Cheon, Bibo Zhu, Ruixuan Zhang, Cori E. Fain, Shengen Shawn Hu, Harish Narasimhan, Gislane de Almeida Santos, Katayoun Ayasoufi, Aaron J. Johnson, Hui Zong, Chongzhi Zang, Haidong Dong, Jie Sun

## Abstract

Aging is a major risk factor for increased morbidity and mortality following acute respiratory virus infections. To elucidate the immune determinants underlying viral pathogenesis and delayed lung repair in the aged lung, a comprehensive time-course study was conducted. Single-cell RNA sequencing (scRNAseq) and high-dimensional flow cytometry were utilized to compare lungs from young and aged mice infected with influenza A virus (IAV). Aged hosts displayed diminished alveolar macrophage (AM) and dendritic cell (DC) but elevated monocyte-derived macrophage (MoM) and interstitial macrophage (IM) presence following infection. Additionally, enhanced accumulation of adaptive immune cells, including CD4^+^ tissue-resident helper (T_RH_) cells, CD8^+^ tissue-resident memory (T_RM_) cells, and a B cell subset resembling age-associated B cells, was observed in the memory phase. Pathway analysis revealed that elevated type I and II interferon (IFNα/γ) signaling, especially in MoM/IM subsets, distinguished the aged hosts from the young. Inhibition of IFNα/γ signaling after viral clearance improved long-term respiratory outcomes and reduced both IM and T_RH_ populations in aged mice. These findings highlight the pivotal role of IFNα/γ signaling, likely within MoM/IM subsets, in driving the exuberant persistence of adaptive immune cells and chronic immunopathology in the aged lung following acute viral infection.

## Introduction

Aging is among the major risk factors for severe morbidity and mortality following acute respiratory infections (Mertz et al. 2013; Goplen et al. 2020). Diminished vaccine efficacy and immune dysregulation further enhance the vulnerability of the elderly to pathogens such as influenza viruses (Silva-Cayetano et al. 2023; Henry et al. 2019). Individuals over 65 years old exhibit increased hospitalization, ICU admissions, and mortality following influenza virus infection (Mertz et al. 2013). Beyond the acute phase, persistent pulmonary pathology, including chronic inflammation, fibrosis and impaired lung function, contributes to a substantial post-acute health burden (Narasimhan et al. 2022; Wei et al. 2023). Recent studies of hospitalized COVID-19 and seasonal influenza patients suggest that ∼50% of respiratory complications (cough, hypoxemia, interstitial lung disease, and shortness of breath) manifest or persist after the acute phase (≥30 days post-admission) (Xie et al. 2024). Concordantly, in murine models, these long-term sequelae were displayed as delayed recovery, excessive collagen deposition and persistent lung inflammation (Narasimhan et al. 2024; Goplen et al. 2020).

While the immune response is essential for viral clearance, maladaptive inflammation and dysregulated immune reconstitution are major drivers of age-related morbidity and mortality (Wu, Goplen, et al. 2021). Multiple layers of immune factors contributed to suboptimal antiviral defenses and tissue damage displayed in aged hosts (Hernandez-Vargas et al. 2014). Relevant aspects include defects in the number and function of alveolar macrophages (AMs) (Wong et al. 2017; McQuattie-Pimentel et al. 2021; Wu et al. 2023), exuberant innate inflammation (Hernandez-Vargas et al. 2014; Kulkarni et al. 2019), reduced T cell receptor (TCR) repertoire diversity (Thome et al. 2016), as well as impaired T and B cell priming (Zhao et al. 2011; Silva-Cayetano et al. 2023). During recovery, persistent inflammation and dysfunctional pro-repair responses aggravate chronic pathology (Goplen et al. 2020; Narasimhan et al. 2024). Both intrinsic (cell-intrinsic aging) and extrinsic (aged tissue environment) factors could account for the unique immune phenotypes that perpetuate lung sequelae.

Interstitial macrophages (IMs) reside in the lung parenchyma and, unlike AMs, are dependent on CSF1 (M-CSF) for development or survival (Aegerter et al. 2022). Their roles in lung homeostasis, tissue repair and disease development are less understood. However, recent studies suggest IMs may serve as a reservoir for viral antigens (Wu et al. 2024), mediate fibrosis (Meziani et al. 2018; Joshi et al. 2020; Aran et al. 2019; Morse et al. 2019; Hoeft et al. 2023), and clear apoptotic alveolar epithelial cells (Zuttion et al. 2024). Certain IM subpopulations have also been implicated in the assembly of inducible bronchus-associated lymphoid tissue (iBALT), hinting that IMs may orchestrate “hubs” for local immunity (Li, Mara, et al. 2024).

To dissect the cellular and molecular mechanisms of influenza-induced chronic lung pathology in aged hosts, we performed time-course single-cell RNA sequencing (scRNAseq) profiling of lungs from young and aged mice up to ∼60 days post infection (d.p.i.). Refined knowledge-based cell type annotation was performed in major immune cell populations, followed by validation with high-dimensional flow cytometry. Persistent inflammation and aberrant macrophage dynamics were observed. During memory phase, aged lungs demonstrated enhanced accumulation of adaptive immune cells, including tissue-resident helper CD4 T cells (T_RH_) (Son et al. 2021; Swarnalekha et al. 2021), CD8 tissue-resident memory cells (T_RM_), and a B cell subset resembling “Age-Associated B cells” (Cancro 2020; Dai et al. 2024). The close association between cell numbers of IMs and these adaptive populations suggests the potential role of IMs in driving their recruitment and/or maintenance. Gene set enrichment analysis (GSEA) further identified elevated interferon (IFNα/γ) signaling, predominantly in lung MoM/IM subsets, in aged hosts. Importantly, therapeutic blockade of IFNα/γ signaling after viral clearance ameliorated chronic lung pathology and reduced IM and T_RH_ accumulation. These findings support a new mechanism in which excessive IFN signaling modulates IMs to promote maladaptive immune niches, offering a promising therapeutic avenue for mitigating long-term lung sequelae in aged individuals.

## Results

### Aging alters immune dynamics and promotes persistent lung inflammation following influenza infection

Aging is a significant risk factor for morbidity and mortality induced by influenza virus infection in both humans (Mertz et al. 2013) and mice (Goplen et al. 2020; Narasimhan et al. 2024). It also influences the dynamics and scale of cellular and molecular changes in the lung after infection (Hernandez-Vargas et al. 2014). This is evident in aged hosts, which showed a delayed onset of the pathogenic process, as demonstrated by initial body weight loss after influenza A H1N1 PR8/34 infection (IAV, PR8) **(Fig. 1A and S1A)** and histological examination of lung tissue stained with H&E **(Fig. 1B)**. Aged hosts also experienced persistent lung inflammation and pathology **(Fig. 1B)**. Investigating the cellular and molecular mechanisms underlying these age-associated alterations requires capturing data at multiple key time points. While previous research has provided detailed characterizations of immune responses during the acute phase (Kasmani et al. 2023), such alterations in aged hosts during the recovery and chronic phases of influenza virus infection are crucial for understanding the pathogenesis of “Long-Flu” observed in aged individuals (Xie et al. 2024).

**Fig. 1.**
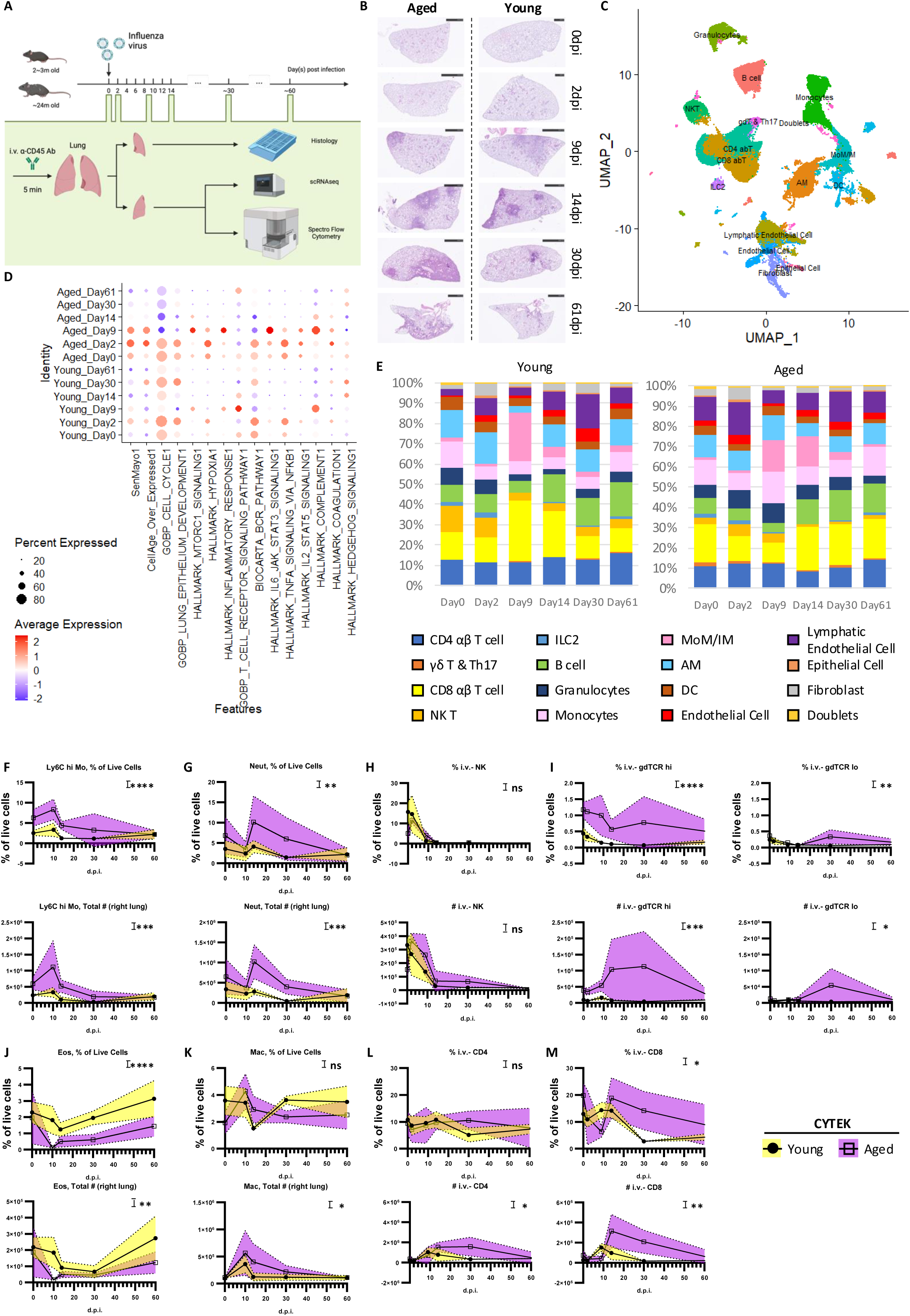
Temporal dynamics of immune cells in the lung of young and aged hosts following influenza virus infection. A. Experimental design for temporal analysis of lung cellular responses in young or aged hosts post influenza virus infection. Young (2-3 months old) and aged (∼24 months old) mice were infected with the same dose of PR8. Samples were collected on 0, 2, 9, 14, 30, and ∼60 days post infection (d.p.i). To label circulating/vasculature-associated immune cells, anti-CD45 antibodies were injected intravenously (i.v.) 5 minutes before euthanasia. The right lung homogenate was used to prepare single-cell suspensions for scRNAseq and high-dimensional flow cytometry (CYTEK), while the left lung was processed for histology. B. Representative H&E staining of the left lung. (C-E). Results were analyzed using all events that passed quality control and pooled from all collected samples (n=3-5 for each sample). C. UMAP visualization of combined scRNA-seq samples. Cells were colored by knowledge based cell type annotation. D. Dot plot displaying module scores of selected GSEA pathways. across all time points comparing lungs from young and aged mice. Dot size and color represents the percentage of cells and average expression level, respectively, in given groups (rows) and pathways (columns). E. Bar graphs showing the proportional cell type composition of time points. The cell types were annotated in (C). Results from (C-E) were analyzed with respect to the number of high-quality cells in each scRNAseq samples. (F-M). Kinetics of immune cells in the lung, quantified by CYTEK, including: Ly6C^hi^ monocytes (F), neutrophils (G), i.v.^-^ NK cells (H), i.v.^-^ γδ T cells (I), eosinophils (J), macrophages (K), i.v.^-^CD4 T cells (L), and i.v.^-^ CD8 T cells (M). Each dot represents the mean value for one sample, with error bars reflecting variability. The statistical analysis was performed using two-way ANOVA to assess whether age is a significant source of variation: ns, p ≥ 0.05; *, p < 0.05; **, p < 0.01; ***, p < 0.001; ****, p < 0.0001. See also Supplementary Fig. 1.

To address this, we conducted a time-course scRNAseq evaluation of cellular and molecular profiles in young and aged mice following influenza virus infection **(Fig. 1A)**. Using naïve mice as controls (Day 0), samples were collected at 2, 9, 14, 30, and 61 days post-infection (d.p.i.). To minimize potential biases introduced by biological variables, such as the batch of mice or virus, young (2-3 months old) and aged (∼24 months old) female mice were simultaneously infected with the same dose of PR8 virus **(Fig. 1A)**. Before tissue harvest, anti-CD45 antibody was injected intravascularly (i.v.) to label circulating and vasculature-associated immune cells. Lung tissue was then collected for both histological analysis and preparation of single-cell suspension. Single-cell suspensions from lung homogenates of all mice within each group were pooled before scRNAseq library preparation, ensuring group representation. Additionally, high-dimensional flow cytometry was employed to gain deeper insights into immune cell composition in each lung.

The libraries were integrated and clustered using Seurat(Stuart et al. 2019) **(Fig. 1C)**, and cluster identities were determined based on their distinguishing features **(Fig. S1B)**. Anchored by clusterProfiler (Wu, Hu, et al. 2021), pathway analysis using Gene Set Enrichment Analysis (GSEA) (Subramanian et al. 2005) revealed differences between young and aged hosts at each time point. A “Module Score” was calculated to evaluate general trends and key signaling pathways identified through GSEA, with results visualized by sample **(Fig. 1D)** and cell type **(Fig. S1C)**.

As a proof of concept, cellular senescence was assessed with publicly available gene sets, SenMayo (Saul et al. 2022) and CellAge (overexpressed) (Chatsirisupachai et al. 2019). Although these datasets highlighted different cellular compartments **(Fig. S1C)**, aged hosts consistently displayed enhanced cellular senescence-related profiles **(Fig. 1D)**. Aged lungs also showed reduced regenerative capacity, evidenced by transcriptional profiles suggesting impaired cell-cycle progression and epithelial development, particularly during the repair and chronic phases **(Fig. 1D)**. Additionally, aged lungs exhibited enhanced pro-inflammatory pathways, though TCR- and BCR-signaling pathways were not notably altered **(Fig. 1D)**. Hedgehog signaling, reported to be anti-fibrotic (Wang et al. 2023), also diminished in aged hosts during the repair and chronic phases **(Fig. 1D)**. Collectively, these findings highlight decreased regenerative capacity, enhanced inflammation and a fibrosis-prone transcriptional profile in aged lungs following influenza virus infection.

Quantification of immune cell dynamics in young and aged hosts was enabled by synergizing results from scRNAseq and flow cytometry. In each scRNAseq library, the proportions of different cell types were calculated in relative to the total cell number passing through quality control **(Fig. 1E)**. Spectral flow cytometry was used to validate the dynamics of both innate and adaptive immune compartments **(Fig. 1E-M; Fig. S1D-G)**. Consistent with pathway analyses **(Fig. 1D)**, aging was associated with an enhanced pro-inflammatory profile. While our time points were suboptimal for capturing the peak innate immune response, particularly in young mice (Zhang et al. 2020), aged hosts exhibited enhanced and persistent elevation of inflammatory cells **(Fig. 1E)**. This was reflected by the accumulation of Ly6C^hi^ monocytes **(Fig. 1E, F; S1D)**, neutrophils **(Fig. 1E, G; S1D)**, and macrophages **(Fig. 1E, K; S1D)**. Aged hosts also displayed increased accumulation of other T cells, including Natural Killer T (NKT) cells **(Fig. 1E, H; S1F)** and γδ T cells **(Fig. 1E, I; S1F)**. Interestingly, these were accompanied by defects in eosinophils **(Fig. 1 E, J; S1D)**. Altered dynamics were also observed among major adaptive immune populations in aged hosts **(Fig. 1E, L-M)**. Both CD4 T cells **(Fig. 1E, L; S1E)** and CD8 T cells **(Fig. 1E, M; S1E)** exhibited delayed but enhanced accumulation in aged lungs. Together, these findings suggest a uncoordinated immune response in aged lungs following influenza virus infection.

### Enhanced accumulation of CD8 T cells with Taa-like features in aged lungs at the memory phase

T cells play a critical role in host antiviral responses following influenza virus infection, but they can also contribute to severe lung immunopathology in both the acute and chronic phases of respiratory infections (Wei et al. 2023; Cheon et al. 2021; Narasimhan et al. 2024). To investigate the dynamics of T cell responses in young and aged hosts post-influenza virus infection, we subset the T cells and performed further characterization **(Fig. 2A & S2A)**. Clusters 12 and 22 were identified as proliferating T cells based on cell cycle analysis **(Fig. 2B)**. Despite the accumulation of both resident CD4 and CD8 T cells in the chronic phase **(Fig. 1K, L)**, aged lungs exhibited a reduced proportion of proliferating αβ T cells **(Fig. S2A)**.

**Fig. 2.**
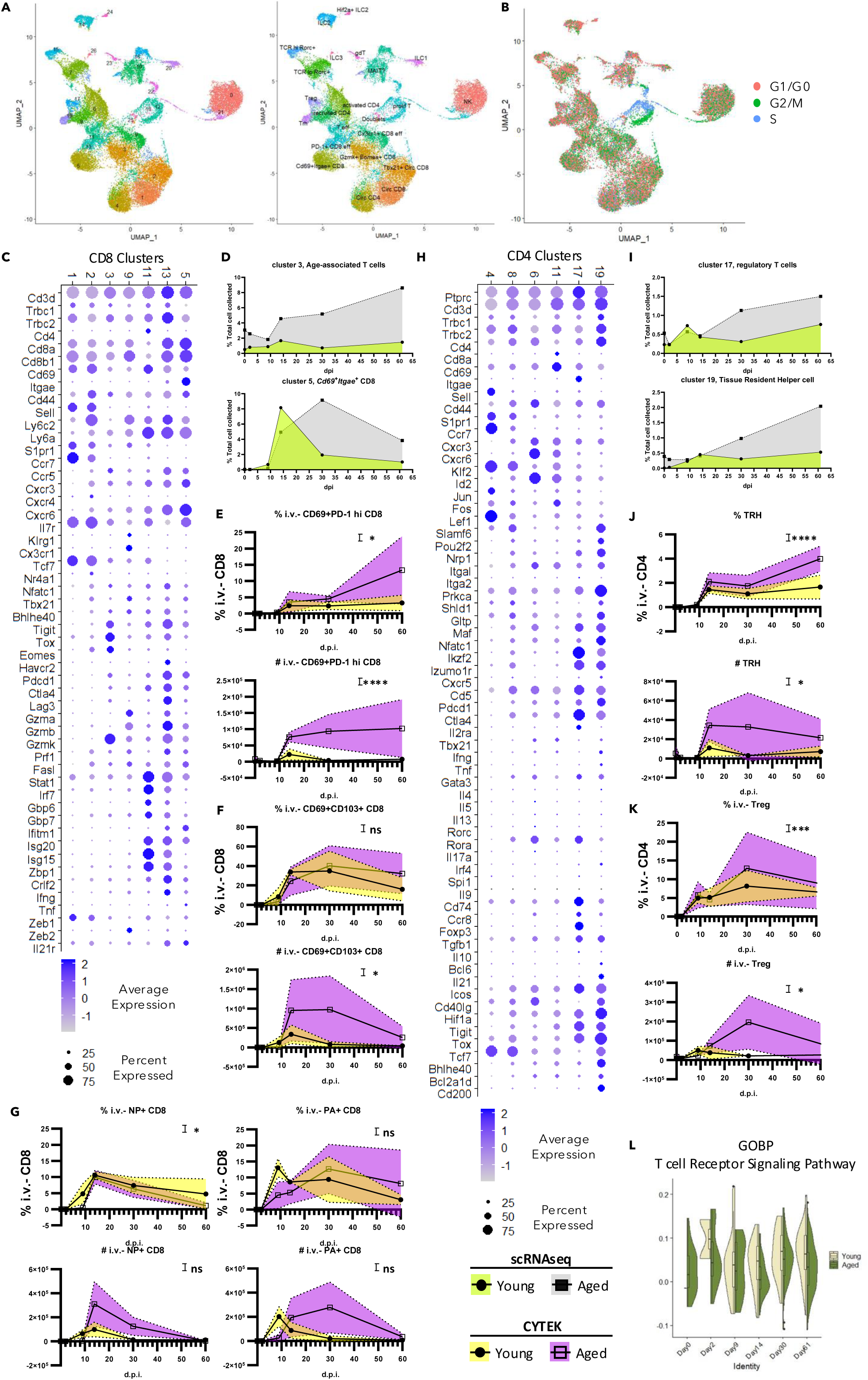
Enhanced T_RM_ and T_RH_ populations are observed in the aged lung during the memory phase. (A-B). UMAP of αβ T cells and invariant T cells identified in Fig. 1C. Cells were colored subsequent re-clustering (A, left), cell sub type annotation (A, right), and cell cycle stage (B). C. Dot plot illustrating the expression level of selected genes in CD8^+^ T cell-related clusters. The color and size of dots represent the average expression and percentage of expressed cells, respectively, for the genes (rows) in clusters (columns). D. Kinetics of age-associated T cells (top) and *Cd69*^+^*Itgae*^+^ T cells (bottom) quantified from scRNAseq. (E-G). Kinetics of CD8^+^ T cell subsets in the lung, quantified by CYTEK, including CD69^+^PD-1^hi^ CD8^+^ T cells (representing age-associated T cells) (E), CD69^+^CD103^+^ CD8^+^ T cells (F), and antigen-specific T cells (G). H. Dot plot showing the expression level of selected genes in CD4^+^ T cell-related clusters. The color and size of dots represent the average expression and percentage of expressed cells, respectively, for the genes (rows) in clusters (columns). I. Kinetics of regulatory T cells (Tregs) (top) and tissue-resident helper T cells (T_RH_) (bottom) quantified from scRNAseq. (J-K). Kinetics of CD4^+^ T cells in the lung, quantified by CYTEK, including TRH (J) and i.v.^-^Tregs (K). L. Violin plot depicting the module score for T cell receptor signaling pathway (MsigDB, MM8546) in TRH cells. The violin shape indicates the distribution density of the scores. The embedded boxplot displays the median (center line), the 25th and 75th percentiles (lower and upper hinges of the box), and whiskers extend to 1.5 times the interquartile range from the hinges. Data in (C, H) are shown as dots. Data were pooled from at least three animals per data point in (E-G; J, K). Each dot represents the mean value for that sample, with color fill indicating error bars. Statistical analysis was performed using two-way ANOVA. The impact of age as a source of variation is indicated as: ns, p ≥ 0.05; *, p < 0.05; **, p < 0.01; ***, p < 0.001; ****, p < 0.0001. See also Supplementary Fig. 2.

Among the CD8 T cells captured, clusters 1 and 2 were identified as circulating CD8 T cells **(Fig. 2C)**. These clusters expressed homing receptors, such as *S1pr1* (S1PR1) and *Ccr7* (CCR7), along with mRNA for central memory-associated markers like *Cd44* (CD44) and *Sell* (CD62L). Cluster 2 is likely to be activated and undergoing tissue recruitment. This notion was supported by its enrichment of transcription factors downstream of TCR signaling (e.g., *Eomes*, *Tbx21*, *Nfactc1*, and *Nr4a1*), cytotoxic function-related molecules (e.g., *Prf1*, *Fasl*, *Gzmm*), and chemokine receptors (e.g., *Cxcr3*, *Cxcr4*, and *Cxcr6*). Despite the lack of naïve circulating CD8 T cells, the proportion of recruited CD8 T cells was similar between young and aged mice at early time points **(Fig. S2B)**.

Age-associated T cells (Taa), characterized by their expression of Granzyme K (*Gzmk*), have been reported to be shared across lymphoid and non-lymphoid organs in aged hosts during physiological aging (Mogilenko et al. 2021). Five clusters (clusters 9, 11, 13, 3, and 5) were discovered to contain *Gzmk*-expressing CD8 αβ T cells **(Fig. 2C)**. Clusters 9, 11, and 13 were likely to be effector CD8 T cells for their molecular profile representing cytotoxic T cells **(Fig. 2C)**. Notably, cluster 9 exhibited high expression of *Zeb2* (ZEB2), a transcription factor critical for terminal differentiation of CD8 T cells (Scott et al. 2018) and essential for cytotoxic lymphocyte fate determination during LCMV infection (Giles et al. 2022). In contrast, clusters 11 and 13 predominantly expressed *Zeb1* (ZEB1), a transcription factor linked to T memory cell formation (Giles et al. 2022). Clusters 11 and 13 displayed enriched IFN-induced transcriptomes and expressed *Zbp1* (ZBP1), a molecule implicated in cell death (Karki et al. 2022). Consistently, both clusters gradually declined over time **(Fig. S2B, E)**. Conversely, cells in cluster 9 underwent contraction but persisted until day 60 in both young and aged mice **(Fig. S2B)**.

Among the *Gzmk*-expressing clusters, cluster 3 displayed the highest level of *Gzmk* and a molecular profile closely resembled the previously described “Taa” population (Mogilenko et al. 2021). Cells in this cluster exhibited a pronounced exhaustion-associated transcriptome **(Fig. 2C)**, including co-inhibitory receptors (e.g., *Pdcd1, Ctla4, Tigit, Lag3*) and exhaustion-related transcription factors (e.g., *Tox* and *Eomes*). They also showed low expression of *Tbx21* (T-bet), *Gzma* (Granzyme A), *Gzmb* (Granzyme B), *Prf1* (Perforin 1), and effector cytokines (*Ifng* and *Tnf*). Notably, aged lungs contained increased numbers of these Taa-like cells even at a steady state **(Fig. 2D)**. Such accumulation was enhanced during the memory phase **(Fig. 2D)**. These findings were further validated using spectral flow cytometry **(Fig. S2C)**. There were 3 different *Cd44*^+^*Sell*^-^ clusters (cluster 3, 5, and 9) identified to accumulate during the memory phase **(Fig. 2C, D; S2B)**. Within the CD69^+^CD103^-^ population, cluster 9 cells primarily exhibited a CD69^+^PD-1^lo^ phenotype, whereas cluster 3 cells were mainly CD69^+^PD-1^hi^ **(Fig. 2C, S2C)**. Consistently, we observed an increased number of i.v.^-^CD69^+^CD103^-^PD-1^hi^ CD8 T cells in aged hosts during the memory phase **(Fig. 2E)**. These results may suggest that *Gzmk*^+^ CD8 T cells with Taa-like features could be a prime candidate for pathogenic CD8 T cells causing chronic lung conditions in aged hosts.

Cluster 5 resembled conventional tissue-resident memory T (T_RM_) cells, as evidenced by its expression of *Cd69* (CD69) and *Itgae* (CD103) **(Fig. 2C)**. Although these cells also expressed several inhibitory receptors (e.g., *Pdcd1, Ctla4, Tigit, Lag3*), they did not show elevated levels of exhaustion-related transcription factors (*Tox*, *Eomes*). While both clusters 3 and 5 had reduced cytotoxicity-related profiles relative to cluster 9, cluster 5 maintained a relatively more robust cytotoxic profile. To gain further insights into the characteristics of these cells in aged hosts, we performed pathway analysis using GSEA **(Fig. S2D)**. Compared to cluster 3, cluster 5 cells were enriched in cell adhesion and MHC-I-dependent TCR activation pathways, and they also displayed an enhanced memory formation signature. In young mice, the number of cluster 5 cells peaked around 14 d.p.i., followed by a contraction phase and maintenance at relatively low levels by ∼60 d.p.i. **(Fig. 2D, 2F, S2C)**. In contrast, in aged mice, the proportion of cluster 5 cells peaked around 30 d.p.i.

In summary, influenza virus infection led to increased levels of both conventional T_RM_ and Taa-like cells in the aged lung, potentially contributing to previously reported chronic lung sequelae (Goplen et al. 2020). Flow cytometry analysis of influenza-specific lung T_RM_ cells using H2D^b^-NP_366-374_ (NP) and H2D^b^-PA_224-233_ (PA) tetramers revealed higher frequencies of influenza-specific CD8 T_RM_ cells in aged lungs compared to young lungs during the memory stage (i.e., at 30 and/or 60 d.p.i.) **(Fig. 2G)**, indicating a potential role of virus antigen and/or aging environment such as increased TGF-β level etc. in their accumulation (Goplen et al. 2020).

### CD4 T cells in aged lungs display altered dynamics, with increased Tregs and T_RH_ populations

In the same analysis **(Fig. 2A)**, the CD4 T cell population consisted of cells from clusters 4, 11, 6, 8, 17, and 19 **(Fig. 2H)**. With close proximity to circulating CD8 T cells in the UMAP space **(Fig. 2A)**, cluster 4 was identified as naïve circulating CD4 T cells **(Fig. 2H)**. These cells were enriched in *Klf2* (KLF2) and its target gene *S1pr1* (S1PR1) (Czesnikiewicz-Guzik et al. 2008), while they had yet to upregulate *Maf* (c-MAF) or *Nfatc1* (NFAT1). Consistent with a previous report(Czesnikiewicz-Guzik et al. 2008), our scRNAseq analysis revealed reduced numbers of naïve CD4 T cells **(Fig. S2E)**. In contrast, the remaining five CD4 T cell clusters displayed profiles of antigen-experienced cells **(Fig. 2H)**, characterized by low expression of *Sell* (CD62L) and high levels of *Cd44*, *Nfatc1* (NFAT1), and *Pdcd1* (PD-1). Notably, cluster 8 possessed a similar transcriptome to cluster 4, suggesting that it too represented circulating CD4 T cells. However, cells in cluster 8 also showed increased *Cxcr3* (CXCR3) expression, a chemokine receptor for tissue recruitment of CD4 T cells.

Both cluster 6 and cluster 11 contained TH1-like CD4 T cells that expressed *Tbx*21 (T-bet) and *Ifng* (IFN-γ). They also showed downregulation of *Ccr7* (CCR7) and upregulation of *Cxcr6* (CXCR6), facilitating their homing to lung tissue. Although cluster 6 primarily comprised CD4 T cells, cluster 11 was likely to include both CD4 and CD8 T cells. Compared to cluster 11, cluster 6 cells demonstrated enhanced expression of *Cd40lg* (CD40L), enabling them to receive co-stimulatory signals from antigen-presenting cells. Additionally, since some cells in cluster 6 were also enriched in transcription factors akin to other helper cell subsets (e.g., *Gata3*, *Rora*), we termed this cluster “activated CD4” **(Fig. 2H, S3E)**.

In line with our previous report (Son et al. 2021), the clusters representing regulatory T cells (Tregs) and tissue-resident helper T cells (T_RH_) located close to each other in UMAP **(Fig. 2A)**. Cluster 17 was defined as Tregs **(Fig. 2H)** due to the expression of *Foxp3* (FOXP3) but not *Bcl6* (BCL6). This cluster was also unique in expressing key immunoregulatory cytokines such as *Il10* (IL-10) and *Tgfb1* (TGF-β). Cluster 19, representing previously reported T_RH_ cells (Swarnalekha et al. 2021; Son et al. 2021), was marked by expression of *Izumo1r* (IZUMO1 receptor), *Pdcd1* (PD-1), the transcription factors like *Bcl6* (BCL6) and *Bhlhe40* (BHLHE40), as well as the cytokine *Il21* (IL-21). Both Treg and T_RH_ cells gradually accumulated over time in young and aged hosts **(Fig. 2I)**. Validation of our scRNAseq results was enabled by spectral flow cytometry, identifying Tregs as FOXP3^+^ CD4 T cells and T_RH_ as FOXP3^-^CD44^+^CD62L^-^CD69^+^PD-1^hi^ CD4 T cells **(Fig. S2G)**. Consistent with the scRNAseq quantification **(Fig. 2I)**, increased numbers of Tregs and T_RH_ cells were observed in aged hosts during the memory phase compared to those of the young **(Fig. 2J, K)**. Despite Treg accumulation, previous studies have shown that Tregs in aged lungs display a dysfunctional phenotype in tissue repair following influenza virus infection, and that they can exhibit a Th1/Th17-like phenotype upon in vitro activation (Morales-Nebreda et al. 2021).

TCR signaling and MHC-II dependency have been implicated in maintaining T_RH_ cells in young hosts on ∼60 d.p.i. (Swarnalekha et al. 2021). However, when we evaluated TCR signaling by a Module Score based on a relevant gene set (GO:0050852), we observed a decreasing trend over 14, 30, and 60 d.p.i. **(Fig. 2K)**. To ensure an adequate number of events from young hosts for comparison, pathway analysis was performed on T_RH_ (cluster 19) cells collected at on Day 30 **(Fig. S2H)**. Compared to young hosts, T_RH_ cells from aged mice displayed an enhanced profile for cell migration and adhesion. Additionally, they showed profiles indicating increased cellular stress and altered lipid metabolism. The functional implications and physiological or pathological consequences of these changes in aged hosts remain to be explored.

### Aged lungs harbor enhanced ABC-like B cell subsets indicative of dysfunctional tissue humoral immunity

Differentially expressed genes were identified specifically within isolated B cell clusters **(Fig. 1C)** to enable refined integration and clustering **(Fig. 3A)**. “Newly recruited” B cells were represented by clusters 0, 2, and 3 **(Fig. 3A, S3A)**. These cells expressed *Cr2* (CD21), and *Fcer2a* (CD23), along with high levels of *Ighd* (IGHD). They expressed homing receptors such as *Sell* (CD62L) and *Ccr7* (CCR7), yet lacked genes typically associated with B cell activation.

**Fig. 3.**
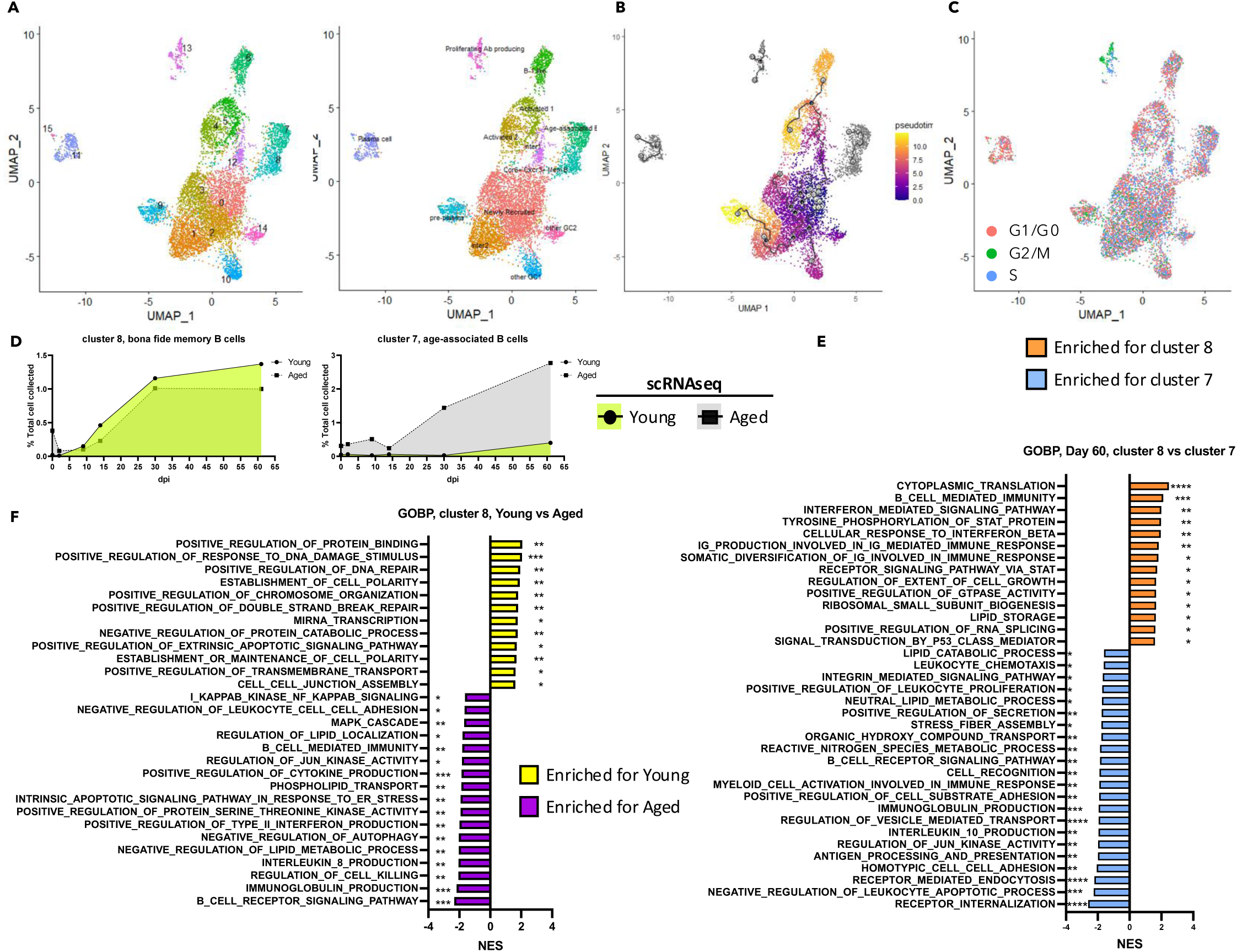
Increased accumulation of class-switched B cells displaying features of age-associated B cell phenotype is observed in aged hosts. (A-C). UMAP of B cells identified in Fig. 1C. Cells were colored subsequent re-clustering (A, left), cell sub type annotation (A, right), cell trajectory (B) and cell cycle stage (C). D. Kinetics of bona fide memory B cells (left) and age-associated B cells (right). E. Bar graph showing normalized enrichment score of selected GSEA pathways generated from ranked differential expressed genes comparing two class-switched B cell populations at 61 days post-infection (d.p.i.). F. Bar graph illustrating normalized enrichment score of selected GSEA pathways generated from ranked differential expressed genes comparing bona fide memory B cells between young and aged hosts across all time points. Data in (E-F) were analyzed with GSEA. Statistical significance is indicated as: *, p < 0.05; **, p < 0.01; ***, p < 0.001; ****, p < 0.0001. See also Supplementary Fig. 3.

However, based on their *Cd44* (CD44) and *Cd38* (CD38) expression, these cells are unlikely to be truly naïve. Throughout the course of infection, aged mice displayed a decreased proportion of these newly recruited B cells **(Fig. S3B)**. Cluster 12 appeared to represent an intermediate stage transitioning from newly recruited to activated B cells **(Fig. 3A-B)**. Meanwhile, clusters 4, 5, and 6 exhibited an activated B cell phenotype preceding class-switch recombination **(Fig. S3A)**. These clusters were characterized by the downregulation of *Ighd* (IGHD) and upregulation of *Nfact1* (NFAT-1), *Nr4a1* (Nur-77), and AP-1 components (*Fos*, *Jun*). They also expressed genes associated with B-1 B cells, including *Zbtb32* (ZBTB32), *Spn* (CD43), and *Cd5* (CD5). Despite their lower proportion of newly recruited B cells, aged lungs contained a higher proportion of activated B cells **(Fig. S3B)**. Interestingly, cluster 6 was almost exclusively detected in aged hosts, peaking around 14 d.p.i.

On a separate route of trajectory **(Fig. 3A-B)**, with cluster 1 (and cluster 2) acting as intermediates, clusters 9, 10, and 14 exhibited germinal center (GC)-like features without evidence of class-switch recombination **(Fig. S3A)**. While lacking the expression of *Igha* (IGHA) and *Ighg1* (IGHG1), these GC-like cells expressed *Cxcr4* (CXCR4), *Cd69* (CD69), *Cd40* (CD40), *Il21r* (IL21R), *Cxcr5* (CXCR5), and *Bcl6* (BCL6). Cluster 9 displayed reduced B cell-related features (e.g., *Cd19*, *Mzb1*). Meanwhile, they had not fully upregulated genes related to plasma cells (*Sdc1*, *Prdm1*, *Irf4*, *Xbp1*). Thus, this cluster was termed as “pre-plasma cell.” Compared to cluster 10, cluster 14 was enriched in genes related to BCR signaling (*Nfact1*, *Nr4a1*) and interferon responses (e.g., *Irf7*, *Isg15*, *Isg20*, *Ifi1*). Unlike clusters 4 and 5, which expressed anti-apoptotic genes (*Bcl2a1a*, *Bcl2a1b*, *Bcl2a1d*), cluster 11 expressed *Zbp1*, a key mediator of cell death under interferon signaling.

Previous results indicated that “immunoglobulin production” and “immunoglobulin receptor binding” were among the top-enriched modules in aged lungs compared to young (Hernandez et al. 2022). Clusters expressed genes of the heavy chains (IgM, IgA, and IgG) included clusters 7, 8, 11, 13, and 15 **(Fig. 3D)**. Both clusters 11 and 15 exhibited transcriptomic profiles resembling plasma cells **(Fig. S3A)**, displaying low levels of B cell markers (*Cd19*, *Mzb1*, *Ebf1*), high levels of plasma cell-associated surface markers (e.g., *Sdc1*, *Ly6c2*) and transcription factors (e.g., *Prdm1*, *Irf4*, *Xbp1*). The combined frequency of these two clusters was elevated in aged hosts **(Fig. S3B)**. Cluster 13 was enriched in *Aicda* (AID), a key enzyme for class-switch recombination (CSR) and somatic hypermutation (SHM) **(Fig. S3A)**. Cell cycle analysis revealed that most cells in this cluster were in S or G2/M phases **(Fig. 3C)**, supporting the notion that they were actively proliferating. These cells largely retained B cell markers **(Fig. S3A)**. Interestingly, while young hosts maintained a robust population of this cluster, aged hosts exhibited a decline in their numbers from 30 dpi to 61 dpi **(Fig. S3B)**.

Cells in clusters 7 and 8 displayed memory B cell-like profiles **(Fig. S3A)**. Cluster 8 cells, which expressed *Ccr6* (CCR6) and *Cxcr3* (CXCR3), resembled previously described bona fide memory B cells that are antigen-specific and confer protection upon secondary challenge (Gregoire et al. 2022). These cells also expressed the costimulatory molecules such as *Icosl* (ICOS-L). While cluster 8 exhibited similar kinetics in both young and aged hosts, cluster 7 cells were notably enriched in aged hosts **(Fig. 3D)**. Beyond pathways associated with cell recruitment and cellular stress, cluster 7 was enriched in pathways linked to innate activation **(Fig. 3E)**. Consistent with this, cluster 7 cells expressed *Tbx21* (T-bet), *Itgax* (CD11c), and *Itgam* (CD11b), along with *Tlr7* (TLR7) and *Tlr9* (TLR9), potentially enabling them to respond to innate stimuli **(Fig. S3A)**. This profile aligns with previously reported “age-associated B cells (ABCs)” that may display autoimmune potential (Nickerson et al. 2023). In contrast, pathways enriched in cluster 8 were related to somatic hypermutation **(Fig. 3E)**. Despite of the similar kinetics between young and aged hosts, the cluster 8 cells in the aged hosts showed defects in DNA repair, as well as diminished cell adhesion and polarity **(Fig. 3F)**. They also exhibited enhanced lipid metabolism, which might fuel increased effector functions.

In summary, both young and aged hosts generate comparable proportions of memory B cells on similar timelines. However, aged hosts appear to have impaired affinity maturation in the lung B cell compartment, potentially explaining reduced vaccine efficacy in senior individuals (Rondy et al. 2017). Additionally, the aged lung harbors a greater abundance of class-switched, antibody-producing B cells with age-associated phenotypes during the memory phase. These cells, possibly activated by innate rather than BCR signaling, may have detrimental effects by producing potential autoreactive antibodies to contribute to chronic diseases after infection.

### Diminished CD11b^+^ PD-L1^-^ DCs in aged mice upon influenza infection

So far, we have demonstrated age-associated accumulation of T_RH_, B cells, and CD8 T cells during the memory phase. However, it remains unclear which cell types orchestrate these responses. Our dataset offers a platform to identify potential candidates. As professional antigen-presenting cells, mononuclear phagocytes (MNPs) are prime candidates for playing such a coordinating role. To dissect their kinetics and functions, we identified clusters containing MNPs and examined their composition and functional states across different time points **(Fig. S4A-S4E)**. Because the scRNAseq library preparation process can skew cell proportions, we also designed a high-dimensional flow cytometry panel using markers suggested by scRNAseq analysis **(Fig. SE)** for validation.

Among the MNPs, dendritic cells (DCs) were identified as cells expressing *Dpp4* (CD26) but not *C5ar1* (CD88) **(Fig. S4C, E)**. We further profiled them using transcription factors, surface markers, and transcriptomic signatures indicative of their functions (Aegerter et al. 2022; Bosteels et al. 2020) **(Fig. S4C)**. Clusters 10 and 29 were characterized as conventional dendritic cells type 1 (cDC1s) for their expression of *Irf8* (IRF8), *Batf3* (BATF3), *Xcr1* (XCR1), and *Itgae* (CD103). Additionally, cluster 29 cells were undergoing proliferation, as revealed by cell cycle analysis **(Fig. S4B)** and their expression of *Mki67* (Ki-67) **(Fig. S4C)**.

Five clusters (17, 21, 27, 30, and 28) were attributed to CD11b^+^ DCs. With proximity to each other on UMAP **(Fig S4A)**, cluster 21 and cluster 17 are likely to be conventional dendritic cells type 2 (cDC2s) **(Fig S4C)**. These clusters were enriched in transcription factors such as *Flt3* (FLT3), *Zbtb46* (ZBTB46), and *Irf4* (IRF4). Notably, cluster 17 expressed *Mgl2* (CD301b), which marks the cDC2s that induce TH-2 response in skin (Kumamoto et al. 2013) and CD8 TRM in the female genital tract (Shin et al. 2016). These cells also expressed *Ccr5* (CCR5), potentially enabling them to respond to chemotactic signals induced by MCP-2, MIP-1α, or MIP-1β.

Compared to cluster 17, cluster 21 expressed *Bex6* (BEX6), previously reported in inflammatory cDC2s during influenza virus infection (4 dpi in young mice) (Bosteels et al. 2020). However, cells in this cluster did not exhibit a fully inflammatory profile like those in cluster 30. Cluster 27 may serve as an intermediate stage between cluster 21 and cluster 30. Clusters 30 and 28 were likely to derive from monocytes, indicated by their expression of *Mafb* (MAFB), *Ccr2* (CCR2), and *Csf1r* (CSF1-R). Positioned close to monocyte-derived macrophages on the UMAP **(Fig. S4C)**, cluster 30 cells expressed high levels of *Irf7* (IRF7) and other interferon-response elements **(Fig. S4C)**. In contrast, cluster 28 lacked inflammation-associated features but was actively proliferating **(Fig. S4B, C)**.

The remaining DC clusters included clusters 20 and 25 **(Fig. S4C)**. Cluster 20 expressed *Flt3* (FLT3) and *Zbtb46* (ZBTB46). Cells in this cluster were also enriched in *Ly75* (CD205), which is important for antigen uptake and processing, and *Ccr7* (CCR7), which is critical for DC migration. These cells also produced chemokines such as *Ccl17* (CCL17) and *Ccl22* (CCL22), potentially facilitating the recruitment of other immune cells including CD4 T cells, NK cells, and γδ T cells. Although limited expression level of *Il12a* (IL12A) was detected, the enrichment of Il12b (IL12B) suggests that these DCs may retain the capacity to produce IL-12, a cytokine crucial for TH1 responses (Heufler et al. 1996) . Positioned apart from other DC clusters in the UMAP **(Fig. S4A)**, cluster 25 was identified as plasmacytoid DCs (pDCs) due to the expression of *Siglech* (Siglec-H), *Ly6c2* (Ly6C), *Ly6d* (Ly6D), and *Tcf4* (TCF-4).

Kinetics of different DC subsets were evaluated with scRNAseq analysis and flow cytometry. Overall, DC kinetics were similar between young and aged hosts **(Fig. S1D, S4F)**. Although the proportions of clusters 10 and 29 (cDC1 subsets) declined in aged hosts compared to young controls **(Fig. S4D)**, no significant difference in the cDC1 population was observed at 30 d.p.i. **(Fig. S4G, H)**. Using PD-L1 as a surface marker, we noted a reduction in the CD11b^+^PD-L1^-^ DC population **(Fig. S4G, I)**, which primarily consists of cDC2 cells (Fig. S4G). Further examination of the scRNAseq data indicated that this decrease was mainly attributable to a reduction in cluster 17 (*Mgl2*^+^ cDC2s) **(Fig. S4D)**. Despite the critical role of conventional DCs in supporting T cells, we found no clear synchronization in their abundance alongside adaptive immune cells.

### Interstitial macrophages as a potential key mediator of age-related adaptive immune cell accumulation

To gain further insight into monocytes and macrophages, we re-clustered these cells **(Fig. S4A, E; Fig. 4A)**. The lung harbors two major types of resident macrophages: Alveolar Macrophages (AMs) and interstitial macrophages (IMs). In our dataset **(Fig. 4A)**, we identified AMs (clusters 12, 18, 11, 19, 4, 0, 14, 24, and 16) using previously reported features (Aegerter et al. 2022) and transcription factors (e.g. *Pparg*, *Bhlhe40*, *Bhlhe41*) known to be important for AM identity and proliferation **(Fig. S5A)**. Because of the previously reported function of *Maf* (c-MAF) and *Mafb* (MAF-B) in inhibiting the embryonic stem cell (ESC)-like features in AMs (Soucie et al. 2016), we used this as an exclusion criterion. Consistent with previous reports (Chakarov et al. 2019), we identified two main IM clusters (13 and 25) using known IM markers (Aegerter et al. 2022) **(Fig. S5A)**. Both clusters expressed *Mafb* (MAF-B) (Vanneste et al. 2023), a transcription factor implicated in IM fate decisions. They also expressed co-stimulatory molecules such as *Cd40* (CD40), *Cd80* (CD80), and *Cd86* (CD86), and expressed *Tgfb1* (TGFβ), a cytokine associated with the establishment of resident T cells (Zhang & Bevan 2013; Wu et al. 2020).

**Fig. 4.**
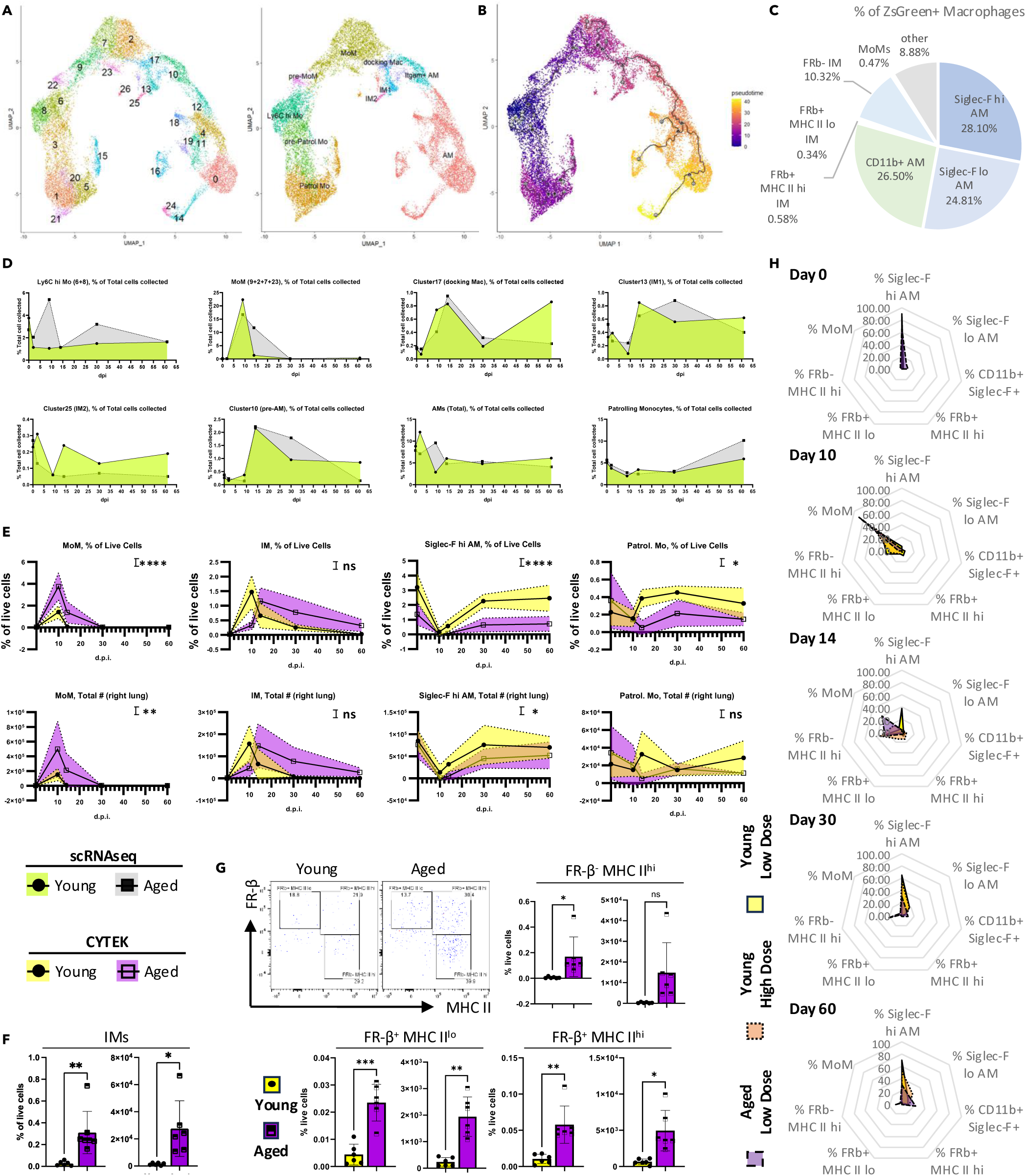
IMs emerge as a promising candidate regulating the accumulation of adaptive immune cells in aged hosts. (A-B). UMAP of monocytes and macrophages identified in Fig. S4A. Cells were colored subsequent re-clustering (A, left), cell sub type annotation (A, right), and cell trajectory (B). C. Lineage tracing using Ccr2^iCre^ ZsGreen^+^ mice. Mice were infected with PR8, followed by daily tamoxifen treatment starting at 13 d.p.i. for 5 days before tissue harvest at 30 d.p.i. The pie chart indicates the composition of ZsGreen^+^ macrophages. D. Kinetics of monocytes/macrophages quantified by scRNAseq (from Fig. 4A) expressed as a proportion of total quality-controlled events presented in Fig. 1C. E. Kinetics of monocytes/macrophages in the lung quantified by high dimensional flow cytometry (CYTEK). Cells examined include monocyte-derived macrophages (MoM), interstitial macrophages (IMs), Siglec-F^hi^ alveolar macrophages (AMs), and patrolling monocytes (Patrol. Mo). F. Bar graph showing IM quantification at ∼60 d.p.i. G. Representative plots (upper left) and bar graphs quantifying IM subsets. H. A group of young mice infected with a higher dose of PR8 (orange) was included along with young (yellow) and aged (purple) mice infected with the same lower dose of PR8. Lungs were harvested at 0, 10, 14, 30, and ∼60 d.p.i. Macrophage composition at each time point is presented in radar plots, where each axis represents a distinct macrophage population. Radar plot axes are arranged in an anticlockwise direction to roughly reflect the inferred differentiation trajectory of macrophages shown in Fig 4B. Data were pooled from at least three animals per data point (E-H). In (E), each dot represents the mean value of a sample, with color-filled error bars. Statistical analysis was performed using two-way ANOVA. In (F), results were combined from two experiments; each dot represents one animal and statistical analysis was performed with an unpaired Student’s t test with Welch’s correction. Data are shown with respect to whether the age of the mice served as a source of variation: ns, p ≥ 0.05; *, p < 0.05; **, p < 0.01; ***, p < 0.001; ****, p < 0.0001. See also Supplementary Fig. 4 and Supplementary Fig. 5.

Looking into their molecular profile **(Fig. S5A)**, cells in cluster 13 showed higher MHC II expression (e.g., *H2-Ab1*, *H2-DMb1*), potentially providing a strong signal 1 for T cell activation. These macrophages also displayed elevated levels of the co-stimulatory molecule like *Icosl* (ICOS-L). Conversely, cluster 25 had relatively low MHC II expression and expressed *Lyve1* (LYVE-1) and *Folr2* (FR-β). Its transcriptome resembled “M2-like” macrophages, marked by *Cd163* (CD163), *Mrc1* (CD206), *Retnla* (RELMα), and *Il10* (IL-10). This cluster displayed features reminiscent of *Pf4*^+^*Mrc1*^hi^ IMs reported in allergy and bacterial infection models (Li, Mara, et al. 2024). Hereafter, we refer to cluster 13 as “IM1” and cluster 25 as “IM2.”

At homeostasis (Day 0), we detected classical/inflammatory monocytes (clusters 8 and 6), nonclassical/patrolling monocytes (clusters 1, 21, 5, 15, 26), and intermediates between these two states (clusters 3, 20) **(Fig. S5B, 4A, D)**. Patrolling monocytes were identified as *Spn* (CD43)^+^, *Cx3cr1* (CX3CR1)^hi^, *Ly6c* (Ly-6C)^-^, and *Ccr2* (CCR2)^-^ cells **(Fig. S5A)**. They also expressed *Nr4a1* (Nur77) and *Pou2f2* (OCT2). Meanwhile, classical/inflammatory monocytes expressed high levels of *Ly6c2* (Ly-6C), along with homing receptors such as *Ccr2* (CCR2) and *Cx3cr1* (CX3CR1). To note, they had not yet upregulated macrophage markers like MHC II or *Mertk* (MERTK). Previous studies have shown that Ly6C^hi^ monocytes can differentiate into both IMs (Vanneste et al. 2023) and monocyte-derived AMs (MoAMs) (Li et al. 2022). Thus, we selected cluster 8 as the starting population for trajectory analysis **(Fig. 4A, B)**.

Upon infection, infiltrating monocytes (clusters 8 and 6) gradually differentiate into monocyte-derived macrophages (MoMs) and subsequently commit to various fates **(Fig. 4B; S5A, B)**. Compared to monocytes, MoMs (clusters 9, 7, 2, and 23) expressed *Fcgr1* (CD64), *Mertk* (MERTK), and mRNAs for MHC II **(Fig. S5A)**. Despite the expression of co-stimulatory molecules (e.g., *Cd40*, *Cd80*, *Cd86*), the pattern for expression levels was quite distinct between MoMs and IMs **(Fig. S5A)**. Clusters 9 and 7 showed heightened interferon signaling and cell death-related components, while clusters 7 and 2 expressed pro-healing markers such as Arg1 (Arginase-1) **(Fig. S5A)**. These findings suggest that MoMs not only serve as major pro-inflammatory myeloid cells during influenza virus infection but also act as a critical node, transitioning toward the repair phase by altering their phenotype and fate. Indeed, lineage tracing using *Ccr2*^iCreZsGreen^ mice confirmed that infiltrating monocytes could give rise to all macrophage subsets **(Fig. 4C)**, and these labeled cells persisted in the lung until 30 d.p.i.

To further illustrate the roles of monocytes and macrophages during influenza infection, we quantified their kinetics using both scRNAseq **(Fig. 4D)** and spectral flow cytometry **(Fig. S1D; S5C-E; 4E-H)**. In aged hosts, we observed a persistent population of MoMs **(Fig. 4D, E)**. However, this did not translate into an increased AM population **(Fig. S5D, E; 4E)**. We found a defect in Mature/homeostatic AMs, identified as Siglec F^hi^ cells **(Fig. 4D, S5C)**. On a separate branch of the trajectory **(Fig. 4B)**, IMs showed different kinetics between young and aged hosts **(Fig. 4D)**. Concurrent with an enhanced adaptive immune compartment, IM numbers were elevated during the memory phase in aged hosts **(Fig. 4F)**. Since the two IM clusters displayed distinct features **(Fig. S5A)**, FRβ was incorporated to distinguish these populations **(Fig. 4G; S4E; S5C)**. By flow cytometry, we could further separate FRβ-expressing IMs based on their MHC II expression **(Fig. 4G**). Aging was associated with an increased accumulation of all these IM populations during the memory phase **(Fig. 5G)**.

**Fig. 5.**
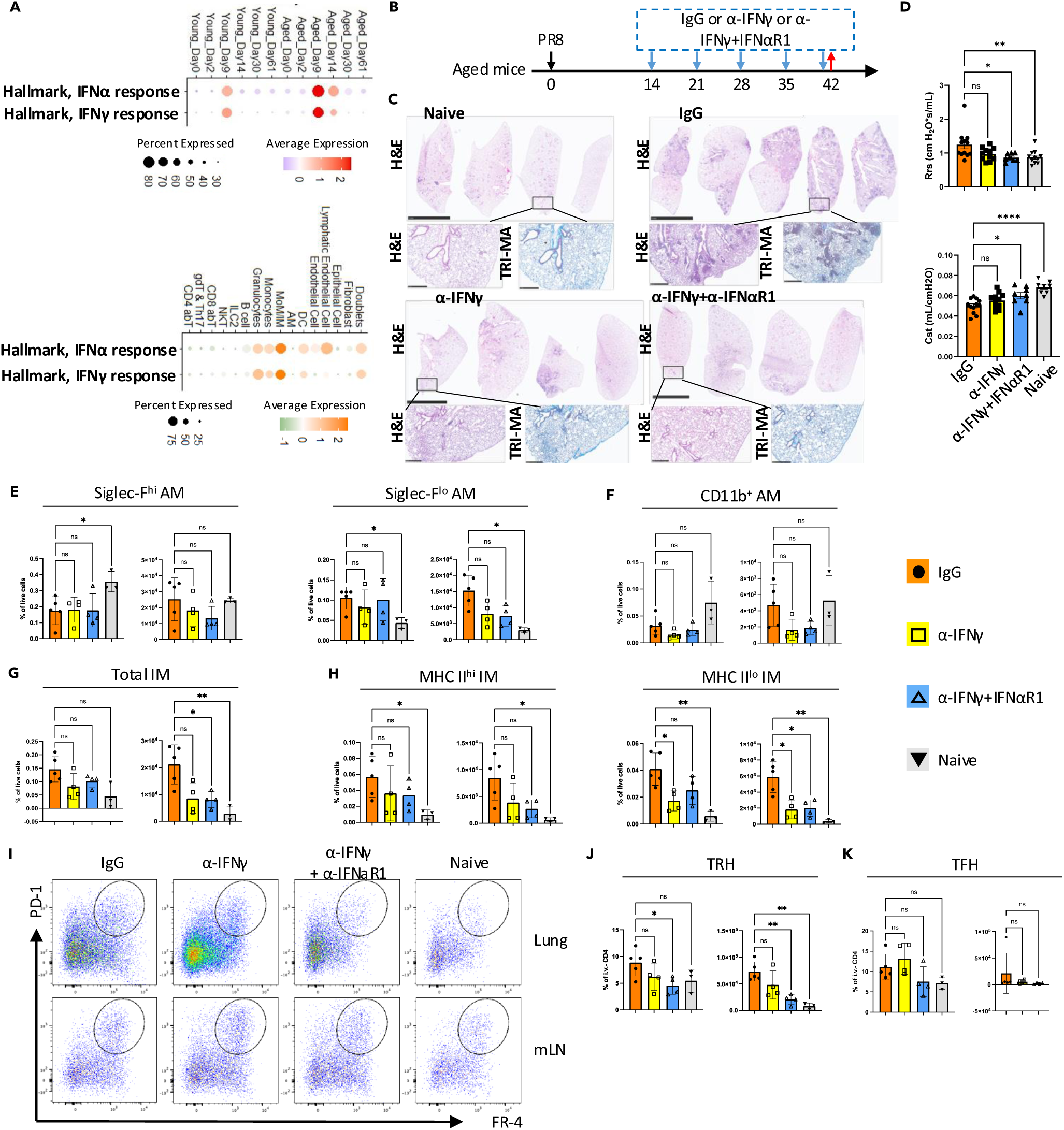
Exuberant type I and type II interferon signaling synergistically drives chronic sequelae in the aged lung. A. Module scores for “Hallmark, IFNα response (MSigDB, MM3877)” and “Hallmark, IFNγ response (MSigDB, MM3878)” across all samples (top) and across all cells (bottom). Dot size and color intensity represents the percentage of cells and average expression level, respectively, in given pathways (rows) and groups/cell types (columns). B. Experimental design. Starting on 14 d.p.i., infected aged mice received weekly treatments of anti-IFNγ ± anti-IFNαR1 monoclonal antibodies (i.p.). Mice were euthanized on 42 d.p.i., with the left lung collected for histology and the right lung for flow cytometry. C. Representative H&E or Masson’s trichrome staining of lung sections from indicated groups. D. Evaluation of resistance (Rrs) and compliance (Cst) of the respiratory system with flexiVent in infected aged mice post indicated treatment. (E-H). Quantification of macrophage populations, including alveolar macrophages (AMs, E-F) and interstitial macrophages (IMs, G-H). I. Representative flow cytometry plots of T_RH_ in the lung and TFH in the mLN for each treatment group. J. Bar graph showing quantification of T_RH_. K. Bar graph showing quantification of T_FH_. Statistical analysis in (D-H, J-K) was performed using repeated measures (RM) one-way ANOVA with Geisser-Greenhouse correction and multiple comparisons. Significance is indicated as *, p < 0.05; **, p < 0.01; ***, p < 0.001; ****, p < 0.0001. See also Supplementary Fig. 6.

Previous report has indicated that enhanced tissue damage during the acute phase can lead to long-term sequelae, even in young hosts (Keeler et al. 2018). Since aging is associated with impaired anti-viral responses (Hernandez-Vargas et al. 2014; Kulkarni et al. 2019), we included a group of young mice infected with a higher dose of PR8 to determine if the macrophage composition differences were solely due to increased tissue damage **(Fig. 4H)**. During the key time point (∼14d.p.i.) when the fate choice of MoMs starts to show, the macrophage composition in the young hosts infected with a higher dose of influenza virus displayed an intermediate phenotype. However, in the chronic phase (30d.pi. and 60d.p.i.), they ultimately exhibit a similar profile as young hosts infected with a low dose of influenza virus. In contrast, aged mice consistently exhibited decreased AM proportions and increased IM populations. Thus, the observed alterations are unlikely to be solely due to enhanced acute-phase damage.

Beyond numerical differences, aging also influenced IM phenotypes **(Fig. S5F, G)**. In young hosts, IMs displayed enhanced cell adhesion, cell-cell interactions, and signal transduction. By contrast, IMs in aged hosts showed impaired responses to growth factors and enhanced effector functions. Notably, IM2 cells in aged hosts exhibited enriched pathways related to antigen presentation and the regulation of adaptive immune responses **(Fig. S5G)**. In summary, IMs in the aged lung may serve as a major contributor to the pathogenesis of long-term sequelae observed in aged hosts.

### Exuberant IFNα/γ signaling impairs lung repair in aged hosts

We have demonstrated that aged hosts exhibit persistent lung pathology following influenza virus infection, accompanied by alterations in both innate and adaptive immune cell populations. To identify potential therapeutic targets, we conducted a comparative analysis of the molecular signaling pathways activated upon influenza virus infection in young and aged hosts. Using GSEA (Subramanian et al. 2005) on all cells **(Fig. 1C)**, we discovered that interferon-related pathways were consistently enriched in aged hosts at multiple time points. With the gene lists of “Hallmark, IFNα response (MSigDB, MM3877)” and “Hallmark, IFNγ response (MSigDB, MM3878)”, an overview of these signaling can be evaluated with the scoring of scRNAseq data **(Fig. 5A)**. Notably, aged hosts displayed both prolonged and enhanced interferon signaling. While no significant differences in total lung cells were observed at 61 d.p.i. due to potential signal or sequencing limitations. Meanwhile, previous Nanostring data have shown enhanced interferon signaling in aged hosts on 60 d.p.i. (Goplen et al. 2020). This finding suggests a potential link between sustained interferon signaling and the chronic pathology observed in aged lungs.

Leveraging the scRNAseq approach, we examined the potential sources (senders) and targets (receivers) of interferon signaling **(Fig. S6A-C)**. CD8 T cells and NKT cells were the primary producers of *Ifng* (IFN-γ) **(Fig. S6A)**. For type I interferons, the major producers included monocyte-derived macrophages/interstitial macrophages (MoM/IM) and stromal cells, such as epithelial cells and fibroblasts **(Fig. S6B)**. Strikingly, MoM/IM rather than the adaptive immune populations were the principal recipients of both interferon signals **(Fig. 5A; S6C)**.

Previous studies indicated that persistent IFNγ signaling could drive chronic fibrotic pathology after SARS-CoV-2 infection (Li, Qian, et al. 2024). Based on our analysis, we hypothesized that IFNα and IFNγ signaling might act synergistically in promoting long-term lung damage. To avoid compromising viral clearance (Hernandez-Vargas et al. 2014), weekly treatment with anti-IFNγ ± anti-IFNαR1 monoclonal antibodies starting at 14 d.p.i. was employed **(Fig. 5B)**. Indeed, blocking both signaling pathways led to reduced chronic inflammatory cell infiltration (H&E staining) and collagen deposition (trichrome staining of collagen) **(Fig. 5C)**. Assessments of respiratory resistance (Rrs) and lung compliance (Cst) by FlexiVent affirmed a significant improvement of airway hyperresponsiveness and lung fibrotic change post IFNγ and IFNα signaling blockade **(Fig. 5D)**. While the alveolar macrophage populations remained unchanged **(Fig. 5E-F)**, we observed a decrease in IMs **(Fig. 5G)**, particularly in the MHC II^lo^ subset **(Fig. 5H)**, which has been reported to reside near the vasculature (Chakarov et al. 2019).

Given the essential role of T_RH_ cells in supporting CD8 T cells and B cells in lung (Son et al. 2021; Swarnalekha et al. 2021), we also examined relevant populations **(Fig. 5I-K; S6D-G)**. Compared to anti-IFNγ treatment alone, combined anti-IFNγ + anti-IFNαR1 treatment further reduced both the proportion and absolute number of T_RH_ cells **(Fig. 5I, J)**. Meanwhile, the follicular helper T (T_FH_) cells in the mediastinal lymph node (mLN) remained unaffected **(Fig. 5I, K)**. Tregs, which share a transcriptional resemblance to T_RH_ cells among T cell subsets **(Fig. 2A, H)**, also decreased in the lung but not in the mLN following treatment **(Fig. S6E-G)**.

Interestingly, potential precursor CD4 T cell populations (clusters 6 and 8) exhibited similar kinetics in young and aged hosts **(Fig. 2A, H; S2E)**, suggesting that, rather than diminished generation or recruitment of T_FH_ cells, the observed reduction in T_RH_ (and possibly Treg) cells may be a localized phenomenon within the lung.

In summary, our findings highlight the synergistic role of interferon signaling in driving long-term pathology in aged hosts following influenza virus infection. We identify MoM/IM populations and T_RH_ cells as likely important players in this process, providing a potential avenue for therapeutic intervention.

### Profiling cell-cell interactions reveals mechanisms underlying chronic sequelae in aged hosts

CellChat (Jin et al. 2021) was employed to infer cell-cell interactions based on ligand-receptor expression in different cell types within each scRNAseq library **(Fig. 1C)**. Comparing young and aged hosts at corresponding time points, we identified unique interactions **(Fig. 6A, S7A)**. For example, Secretoglobin 3A2 (SCGB3A2) has been implicated in protecting lung tissue from cigarette smoke-induced damage (Kurotani et al. 2023). As a proof of concept, we detected “UGRP Signaling,” an interaction between *Scg3a2* and *Marco*, specifically in young hosts at 61 d.p.i. **(Fig. 6A, B)**. Meanwhile, IL-17 signaling emerged exclusively in aged hosts at 14 d.p.i. and 30 d.p.i. **(Fig. 6A, C)**. Notably, the *Nrxn3*-*Nlgn2* interaction between IL-17-producing cells **(Fig. 1C, S1B)** and fibroblasts persisted until 61 d.p.i. **(Fig. 6A, D)**. Among the clusters expressing *Rorc* (Rorγt), the *Il17a* (IL-17A)-expressing clusters (15 and 23) increased in proportion during the memory phase **(Fig. 2A; S7B, C)**. This is consistent with the flow cytometry data showing elevated γδ T cells in the aged lung **(Fig. 1H)**. Together with our GSEA results, these findings suggest that aged hosts develop a TH1/TH17-like profile following influenza virus infection, compared to the young hosts.

**Fig. 6.**
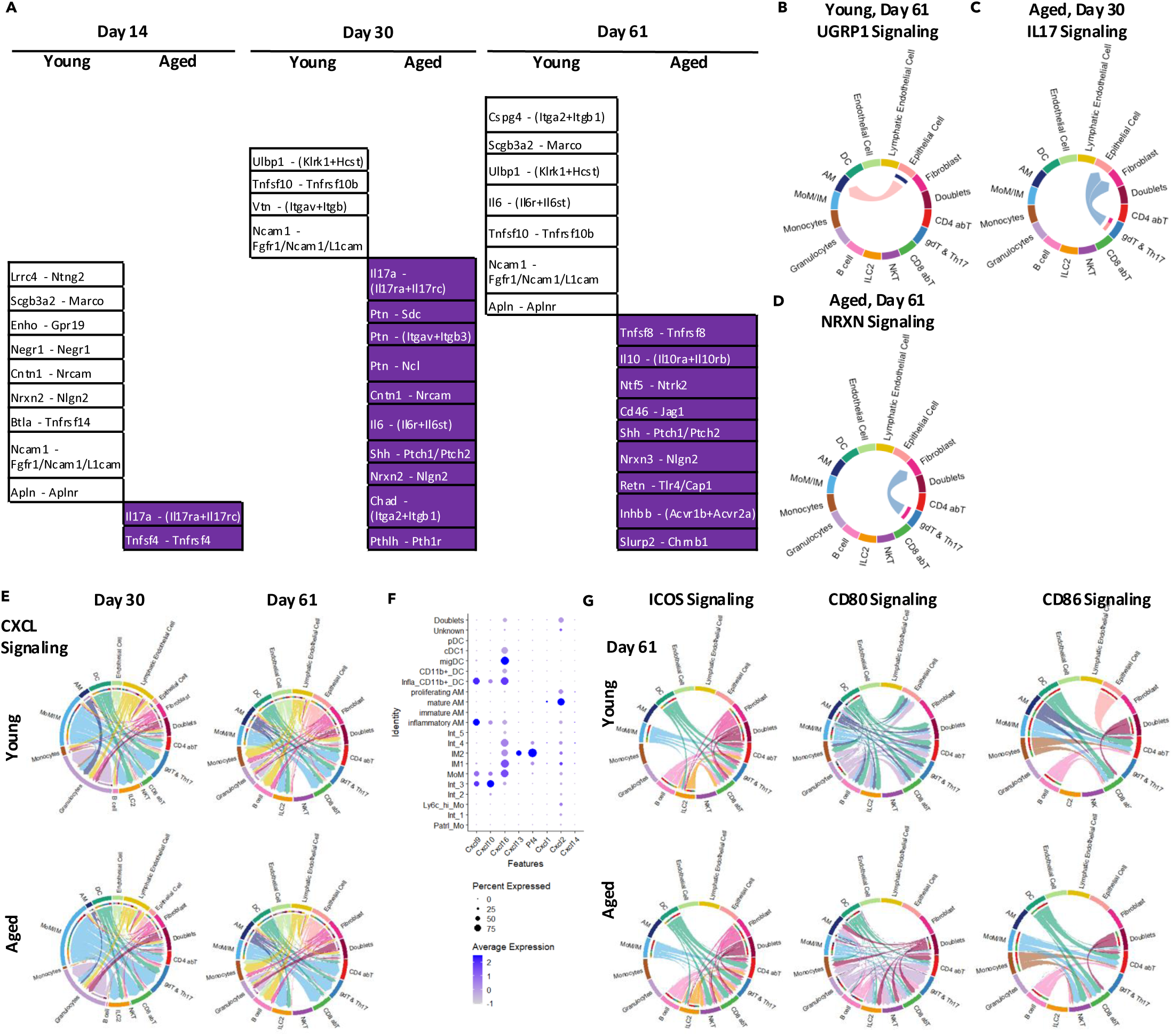
Cell-cell interaction analyses reveal unique pathways in young and aged hosts, as well as potential interactions between IMs and CD4^+^ T cells. A. Unique interactions in young or aged hosts at 14, 30, and 61 d.p.i. (B-D). Chord plot illustrating UGRP1 signaling (Scgb3a2-Marco) in young hosts at 61 d.p.i. (B), IL-17 signaling (Il17a-(Il17ra+Il17rc)) in aged hosts at 30 d.p.i. (C), NRXN signaling (Nrxn3-Nlgn2) in aged hosts at 61 d.p.i. (D). E. Chord plots displaying CXCL signaling in young and aged hosts during the memory phase. F. Dot plot showing ligands involved in CXCL signaling within mononuclear phagocytes (MNPs) (Fig. S4A, C). Dot size and color intensity represents the percentage of cells and average expression level, respectively, in given cell types (rows) and genes (columns). G. Chord plots displaying co-stimulatory signaling at 61 d.p.i. In the chord plots (B-E, G), each arrow indicates an inferred ligand-receptor interaction. Each arrow originates from the signal-sending cell type and points toward the signal-receiving cell type. Arrow color represents the identity of the signal-sending cell, while arrow thickness reflects the inferred interaction strength. See also Supplementary Fig. 7.

Further analysis elucidated interactions between MoM/IM populations and adaptive immune cells. Although some interactions were present in both young and aged hosts, MoM/IMs served as the primary source of “CXCL Signaling” **(Fig. 6E)**. In line with a previous report (Li, Mara, et al. 2024), *Pf4*^+^ IMs (IM2) predominantly produced *Cxcl13* (CXCL13) **(Fig. 6F)**. CXCL13 is the ligand for CXCR5, a chemokine receptor expressed on both T_RH_ (Son et al. 2021) and B cells (Denton et al. 2019). Additionally, cells in the IM2 cluster contributed ligands for “CCL Signaling” **(Fig. S7D-E)**. Besides antigen presentation and chemokine production, MoM/IM cells may also assist adaptive immune cells through co-stimulatory molecules **(Fig. 6G; S4A)**. Collectively, the enhanced accumulation of adaptive immune cells in aged hosts could be linked to increased IM populations through multiple molecular mechanisms.

## Discussion

Upon acute respiratory virus infection, aged hosts display impaired anti-viral responses, heightened inflammation, delayed tissue repair, and an increased risk of chronic lung complications (Wu, Goplen, et al. 2021). Leveraging time-course scRNAseq and high-dimensional flow cytometry, an overview of uncoordinated immune response in aged mice was degranulated. Zooming into the memory phase of the aged lung, enhanced accumulation of adaptive immune cells, particularly CD8 T_RM_, CD4 T_RH_ cells, and a B cell population resembling ABCs (Nickerson et al. 2023; Cancro 2020), were observed. Among myeloid cells, IMs emerged quantitatively as potential key mediators of these accumulations. Consistently, type I and type II interferon (IFN) signaling was enriched in MoM/IM populations, contributing to chronic sequelae. Blocking interferon signaling reduced both IMs and T_RH_ cells, implicating their interactions in sustaining pathogenic immune landscapes. Moreover, the potential assistance from these IMs includes the secretion of chemokines and the ability to provide co-stimulatory signaling. Collectively, we described the synergistic effect of chronic type I and type II interferon signaling in driving tissue immune pathology likely centered around IMs and altered T cell responses in aged hosts.

Through systematic time-course characterization, we have identified several alterations in aged hosts that likely contribute to the pathogenic process in lung upon influenza virus infection. Earlier studies have shown that aged individuals suffer from excessive inflammatory infiltration despite clearance of active virus (Mertz et al. 2013; Hernandez-Vargas et al. 2014; Kulkarni et al. 2019; Goplen et al. 2020). Indeed, the delayed but prolonged presence of inflammatory cells (i.e. neutrophils and Ly6C^hi^ monocytes) can exacerbate tissue injury and hinder regeneration in aged lungs. Moreover, the aged lung environment shifted toward a TH1/TH17-like state, as evidenced by the kinetic analysis of ψοT cells, as well as data with GSEA and CellChat. Lastly, aged lungs harbored both immune cells displaying a dysfunctional phenotype (i.e. AMs) (Wong et al. 2017; Wu et al. 2023) and the ones described as distinct “age-associated” immune subsets (Mogilenko et al. 2021; Nickerson et al. 2023; Cancro 2020; Dai et al. 2024).

Despite expanded memory-like populations including T_RH_ and T_RM_, aged hosts exhibit reduced vaccine efficacy and poorer outcomes after secondary challenges (Goplen et al. 2020). This suggests that these “memory” cells may be functionally compromised upon viral rechallenge. In fact, they might even have detrimental effects: persistent increased T_RH_ (this paper) and T_RM_ responses (Goplen et al. 2020; Narasimhan et al. 2024) correlated with persistent tissue pathology. T_RH_ cells, localized in inducible bronchus-associated lymphoid tissue (iBALT) and capable of assisting both B and CD8 T cells (Son et al. 2021; Swarnalekha et al. 2021), appear central to this phenomenon. Maintenance of T_RH_ cells is thought to be MHC II-dependent (Swarnalekha et al. 2021), raising the question of which local antigen-presenting cells (APCs) sustain them.

Our data pointed to IM populations, especially *Folr2*^+^ IMs, as candidate local APCs that provide both antigenic stimulation and chemokine-mediated recruitment signals, thereby supporting T_RH_ accumulation. Quantitative synchronization has been observed between IMs and T_RH_ upon influenza virus infection. Previous evidence suggested that CD11c^+^ IMs, but not AMs, could phagocytize damaged alveolar type II (AT II) cells during influenza virus infection (Zuttion et al. 2024), enabling the possibility for them to present antigens even after virus clearance. These IMs also express *Cxcl13* (CXCL13), a ligand for CXCR5, allowing chemotactic support for T_RH_ and B cells. Type I interferon signaling, previously shown to induce CXCL13 and promote iBALT formation during acute infection (Denton et al. 2019), may similarly drive pathological memory niches in aged hosts. Blocking IFNγ and IFNαR1 signaling together greatly reduced T_RH_ numbers in the aged lungs but not T_FH_ numbers in mediastinal lymph nodes. This phenomenon highlights a lung-specific mechanism in sustaining T_RH_ responses. Such interplay between IMs and T_RH_ cells *in situ* could initiate early in infection and persist into the memory phase, reinforcing maladaptive immune profiles.

Type I and II interferons are double-edged swords: while integral to anti-viral defense (Seok Lee et al. 2020; Zhang et al. 2022), excessive interferon activity is linked to tissue damage and chronic disease(Sposito et al. 2021). Our data suggest that the blockade of IFNγ and IFNαR1 could synergistically alleviate chronic pathology. Such phenomenon underscores the pivotal role of interferon-driven IM dysregulation. Intriguingly, IM subpopulations can mirror both M1- and M2-like features, suggesting they transcend the classical polarization theory of macrophages. Rather than broad depletion, modulation of IM functions could be optimized aiming for beneficial tissue repair mechanisms while minimizing fibrotic or inflammatory sequelae (Chakarov et al. 2019; Li, Mara, et al. 2024; Goplen et al. 2020).

T cell-intrinsic aging factors could further shape these dynamics. Alterations in T cell transcriptional regulators (e.g., HELIOS/*Ikzf2*) can impose a tissue-infiltration-prone and effector-skewing effect and aged CD4 T cells (Zhang et al. 2023). Although local APC support is likely critical, the mis-location of T_FH_ in the lymph nodes of aged hosts could predispose to alterations caused by insufficient germinal center responses (Jo et al. 2023). Additionally, age-related changes in T cell quality, TCR specificity and bystander activation (Gerlach et al. 2016; Lee et al. 2022) may also influence T_RH_ accumulation and function. Addressing antigen specificity, TCR clonotypes, and spatial organization could further elucidate how T_RH_ and IM interactions reshape local immunity in aging hosts.

In summary, our kinetical and integrated analysis highlights a complex interplay between local immune cells (IMs and T cells) and anti-viral pathways (interferon signaling) in shaping persistent pathology in aged lungs post-influenza infection. Philosophically, the phenomenon we observed could represent a compromised mechanism for aged hosts to cope with their declining protective immunity and regenerative capacity as they strive to memorize prior viral insults. It is crucial to further elucidate the cellular and molecular mechanisms relevant to this phenomenon, allowing the possibility of walking the fine line between protective immune response and pathogenic immune response.

## Acknowledgement

Schematic in the manuscript were created with BioRender.com. We thank UVA Flow Cytometry Research Histology Core Facilities, Biorepository and Tissue Research Facility and NIH tetramer core facility for assistance and reagents. The study was in part supported by the US National Institutes of Health grants AI147394, AG069264, AG090337, AG090337, HL170961 and AI176171 to J.S, American Lung Association Catalyst Award to I.S.C. and American Thoracic Society Unrestricted Research Grant to C.L.

## Data availability statement

The single-cell RNA-seq data can be access through GSE271578 on the Gene Expression Omnibus (GEO).

## Supplementary Figures

**Supplementary Fig. 1.**
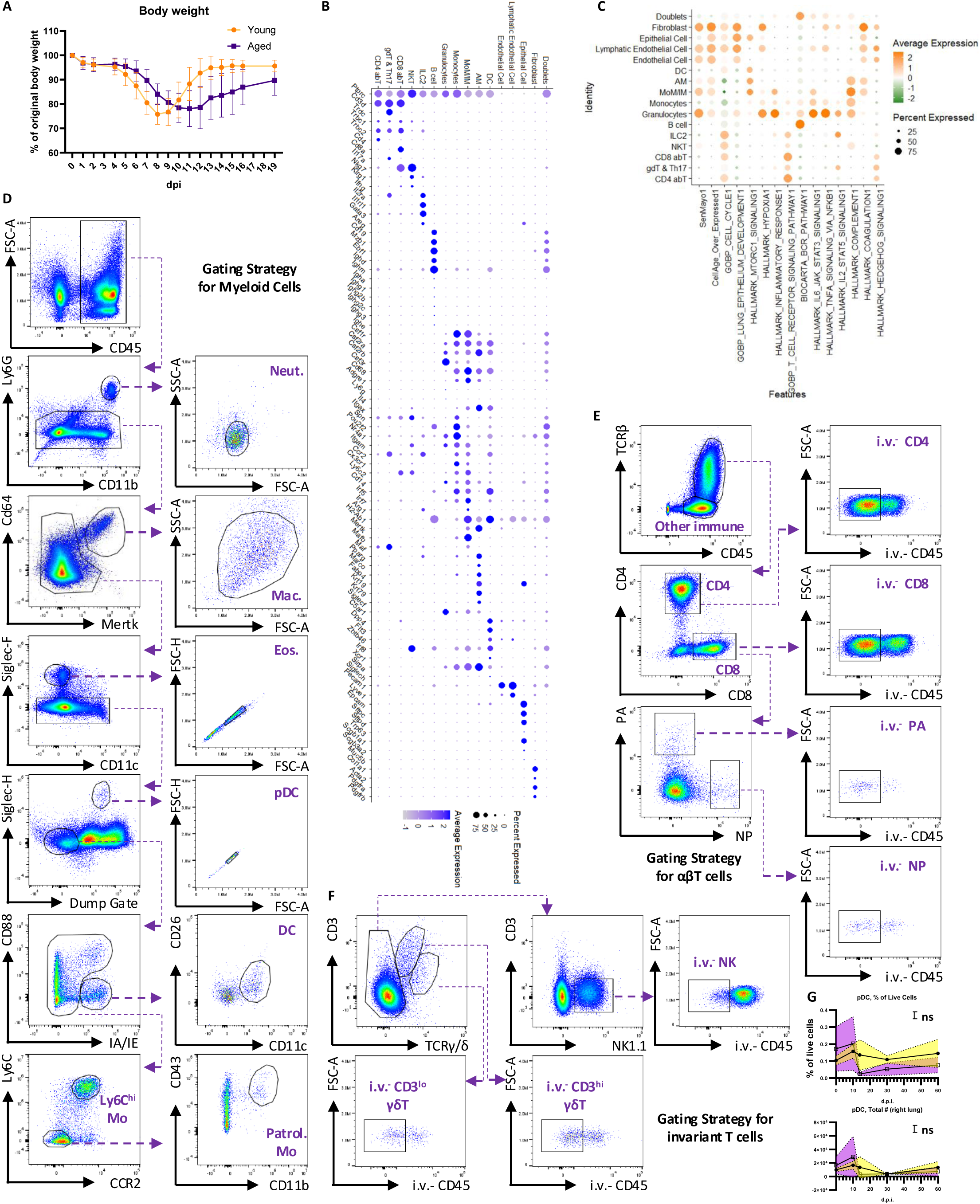
Characterization and immune profiling of the lung in young and aged hosts following influenza virus infection. Young (2-3 months old) and aged (∼24 months old) mice were infected with the same dose of mouse-adapted influenza virus A/PR/8/34 (PR8). Samples were collected at 0, 2, 9, 14, 30, and ∼60 d.p.i. A. Body weight loss curve. B. Features used to define the general immune populations shown in Fig. 1C. Dot size and color intensity represents the percentage of cells and average expression level, respectively, in given genes (rows) and cell types (columns). C. Module scores for selected pathways, displayed as a dot plot for the cell types defined in Fig. 1C. Dot size and color represents the percentage of cells and average expression level, respectively, in given cell types (rows) and pathways (columns). D-F. Gating strategies for myeloid cells (D), αβ T cells (E), and invariant T cells (F) used in spectral flow cytometry. G. Kinetics of plasmacytoid dendritic cells (pDCs) in the lung, quantified by spectral flow cytometry. Data in (G) are pooled from at least three animals per data point. Each dot represents the mean value for that sample, with error bars filled in color. Statistical analysis was performed using two-way ANOVA. Results are shown to indicate whether the age of the mice served as a source of variation: ns, p ≥ 0.05; *, p < 0.05; **, p < 0.01; ***, p < 0.001; ****, p < 0.0001.

**Supplementary Fig. 2.**
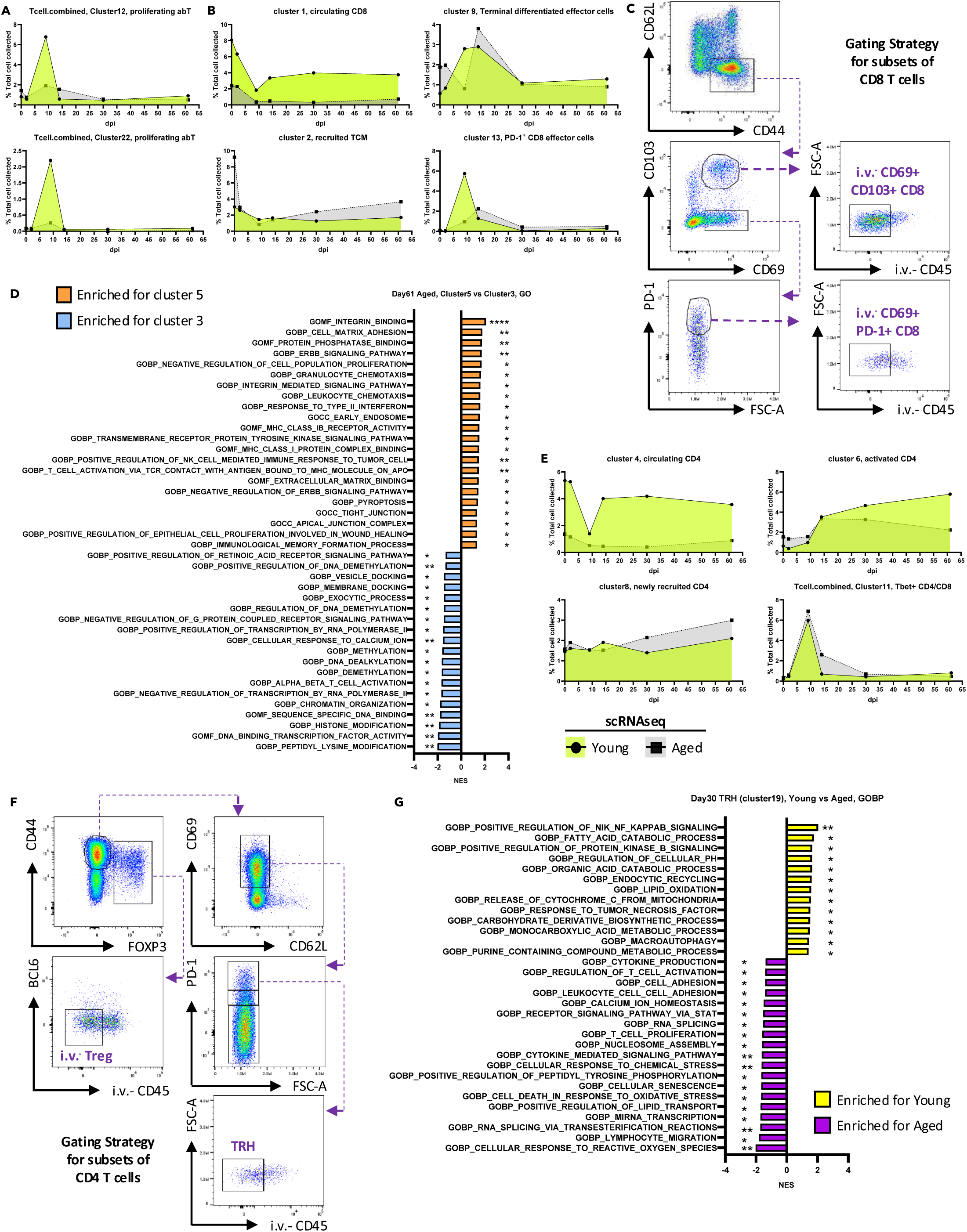
Quantification of αβ T cells using scRNAseq and high-dimensional flow cytometry. A. Kinetics of proliferating αβ T cells quantified from scRNAseq data (Fig. 2A), expressed as a proportion of all events passing quality control in Fig. 1C. B. Kinetics of CD8+ αβ T cells quantified from scRNAseq (Fig. 2A), expressed as a proportion of all quality-controlled events in Fig. 1C. C. Gating strategy for identifying CD69^+^PD-1^hi^ CD8^+^ T cells (representing age-associated T cells) and CD69^+^CD103^+^CD8^+^ T cells by spectral flow cytometry. D. Bar graph showing normalized enrichment score of selected GSEA pathways generated from ranked differential expressed genes comparing two T_RM_ clusters at 61 d.p.i. E. Kinetics of CD4^+^ αβ T cells quantified from scRNAseq (Fig. 2A), expressed as a proportion of all quality-controlled events in Fig. 1C. F. Gating strategy for identifying T_RH_ cells and i.v.^-^ Treg among CD4^+^ T cells by spectral flow cytometry. G. Bar graph showing normalized enrichment score of selected GSEA pathways generated from ranked differential expressed genes comparing TRH populations between young and aged hosts at 30 d.p.i. Data in (D) and (G) were analyzed by GSEA, with significance indicated as *, p < 0.05; **, p < 0.01; ***, p < 0.001; ****, p < 0.0001.

**Supplementary Fig. 3.**
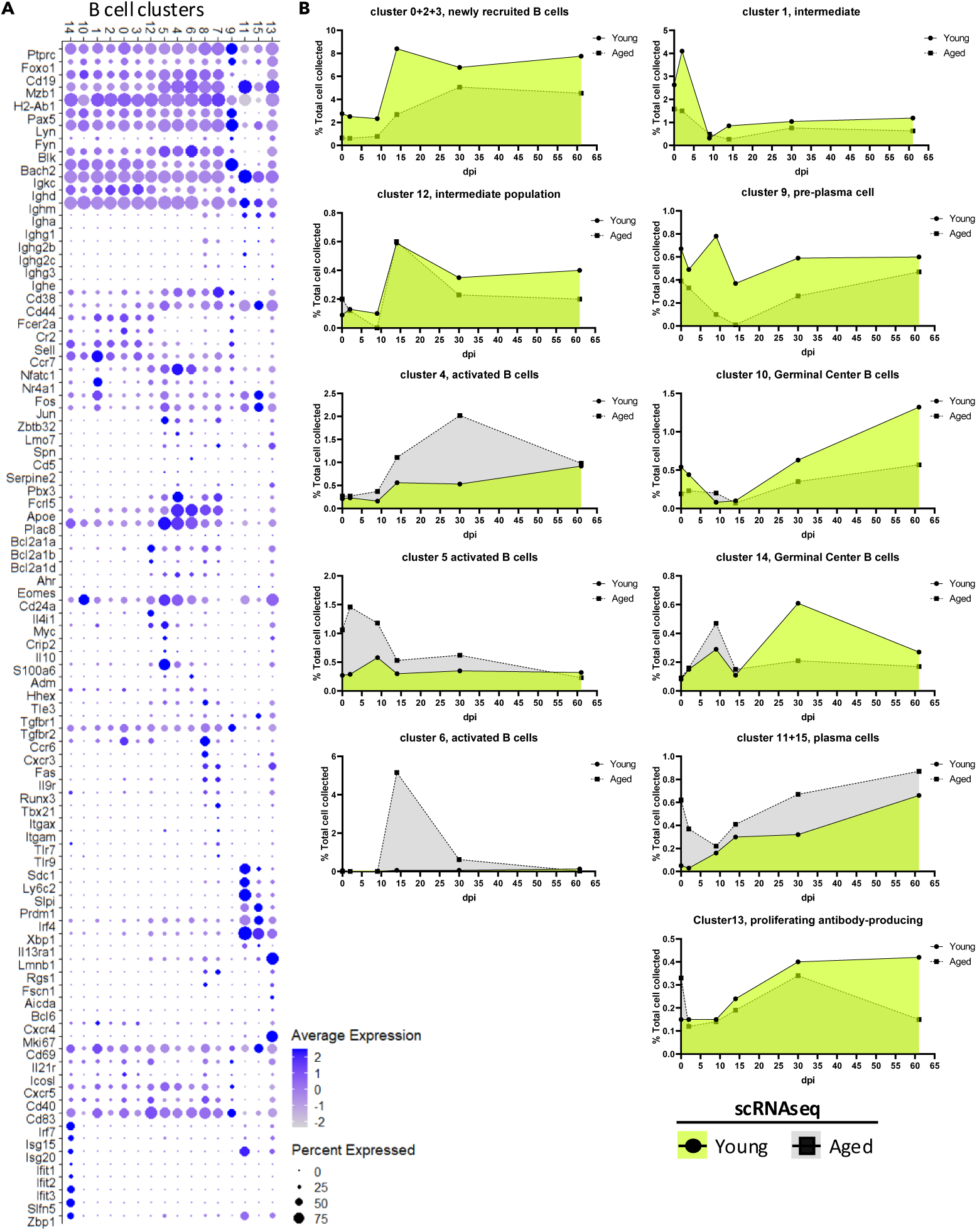
Characterization of B cell clusters in young and aged hosts. A. Dot plot illustrating the defining features of each B cell cluster. B. Kinetics of B cells quantified from the scRNAseq data presented in Fig. 3A, expressed as a proportion of all events passing quality control in Fig. 1C. In (B), each dot’s size represents the percentage of events in each sample expressing the indicated gene, and the color intensity reflects the average expression level.

**Supplementary Fig. 4.**
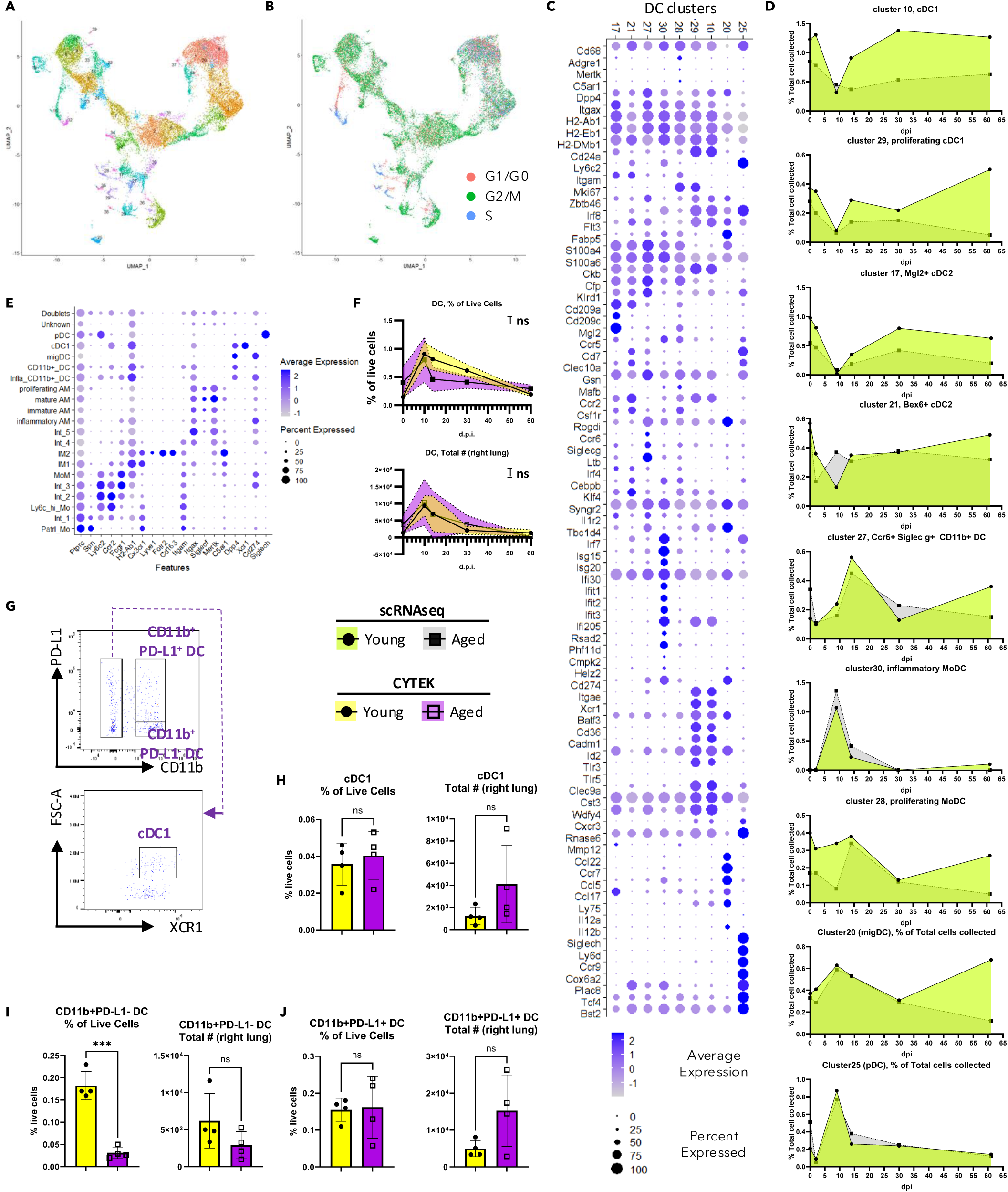
Characterization of mononuclear phagocytes in young and aged hosts. (A-B). UMAP of Mononuclear phagocyte (MNP) identified in Fig. 1C. Cells were colored subsequent re-clustering (A), and cell cycle stage (B). C. Dot plot showing selected features used to define dendritic cell (DC) clusters. Dot size and color intensity represents the percentage of cells and average expression level, respectively, in given genes (rows) and cluster (columns). D. Kinetics of DCs quantified from the scRNAseq data presented in (A), expressed as a proportion of the total quality-controlled events in Fig. 1C. E. Dot plot illustrating selected features used to characterize MNPs by flow cytometry. Dot size and color intensity represents the percentage of cells and average expression level, respectively, in given cell types (rows) and genes (columns). F. Kinetics of DCs (gating strategy shown in Fig. S1D) in the lung, quantified by flow cytometry. G. Gating strategy for DC populations. (H-J). Bar graphs displaying the quantification of DC subsets at ∼60 d.p.i., including cDC1 (H), CD11b^+^PD-L1^-^ DCs (I), and CD11b^+^PD-L1^+^ DCs (J). Data were pooled from at least three animals per time point (F-J). In (F), each dot represents the mean value for a sample, with color-filled error bars. Statistical analysis was performed using two-way ANOVA. In (H-J), results were pooled from two experiments; each dot represents one animal, and statistical analysis was performed using an unpaired Student’s t test with Welch’s correction. Significance levels regarding the impact of age as a source of variation are shown as: ns, p ≥ 0.05; *, p < 0.05; **, p < 0.01; ***, p < 0.001; ****, p < 0.0001.

**Supplementary Fig. 5.**
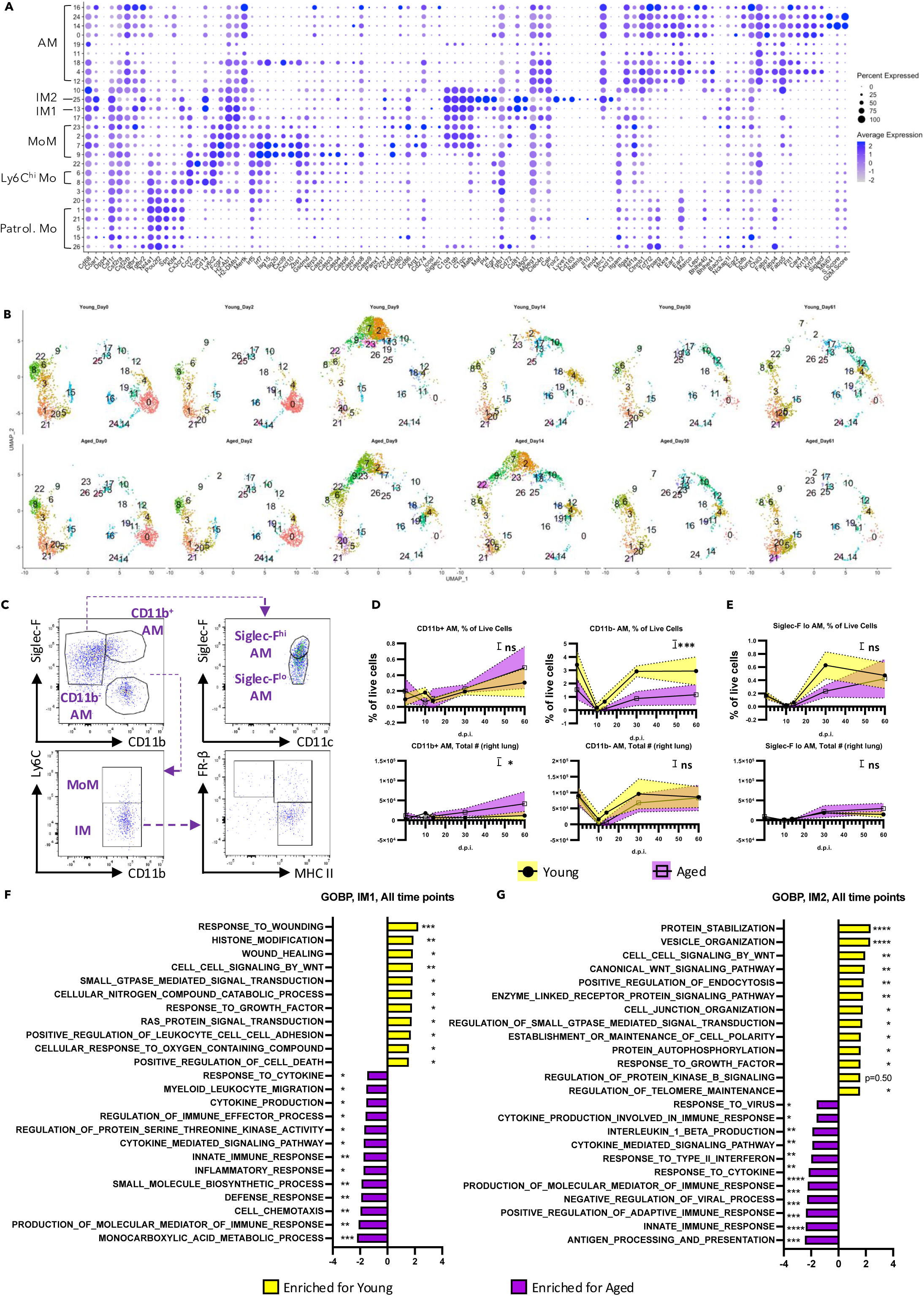
Characterization of monocytes and macrophages in young and aged hosts. A. Dot plot displaying selected features used to define subsets of monocytes and macrophages. Dot size and color intensity represents the percentage of cells and average expression level, respectively, in given clusters (rows) and genes (columns). B. UMAP plots showing the distribution of monocyte/macrophage clusters across with respect to individual samples. C. Gating strategy for identifying macrophage populations by flow cytometry. (D-E). Kinetics of alveolar macrophages (AMs) in the lung, quantified by CYTEK. AM subsets include CD11b^+^ AMs (D, left) and CD11b^-^ AMs (D, right). Within the CD11b^-^ AM population, Siglec-F^hi^ (as shown in Fig. 4E) and Siglec-F^lo^ AMs were quantified (E). (F-G). Bar graphs showing normalized enrichment score of selected GSEA pathways generated from ranked differential expressed genes comparing comparing IM1 (F) or IM2 (G) between young and aged hosts. Data were pooled from at least three animals per data point (D-E). Each dot represents the mean value per sample, with error bars filled in color. Statistical analysis was performed using two-way ANOVA. Data in (F-G) were analyzed by GSEA. Significance levels are indicated as: ns, p ≥ 0.05; *, p < 0.05; **, p < 0.01; ***, p < 0.001; ****, p < 0.0001.

**Supplementary Fig. 6.**
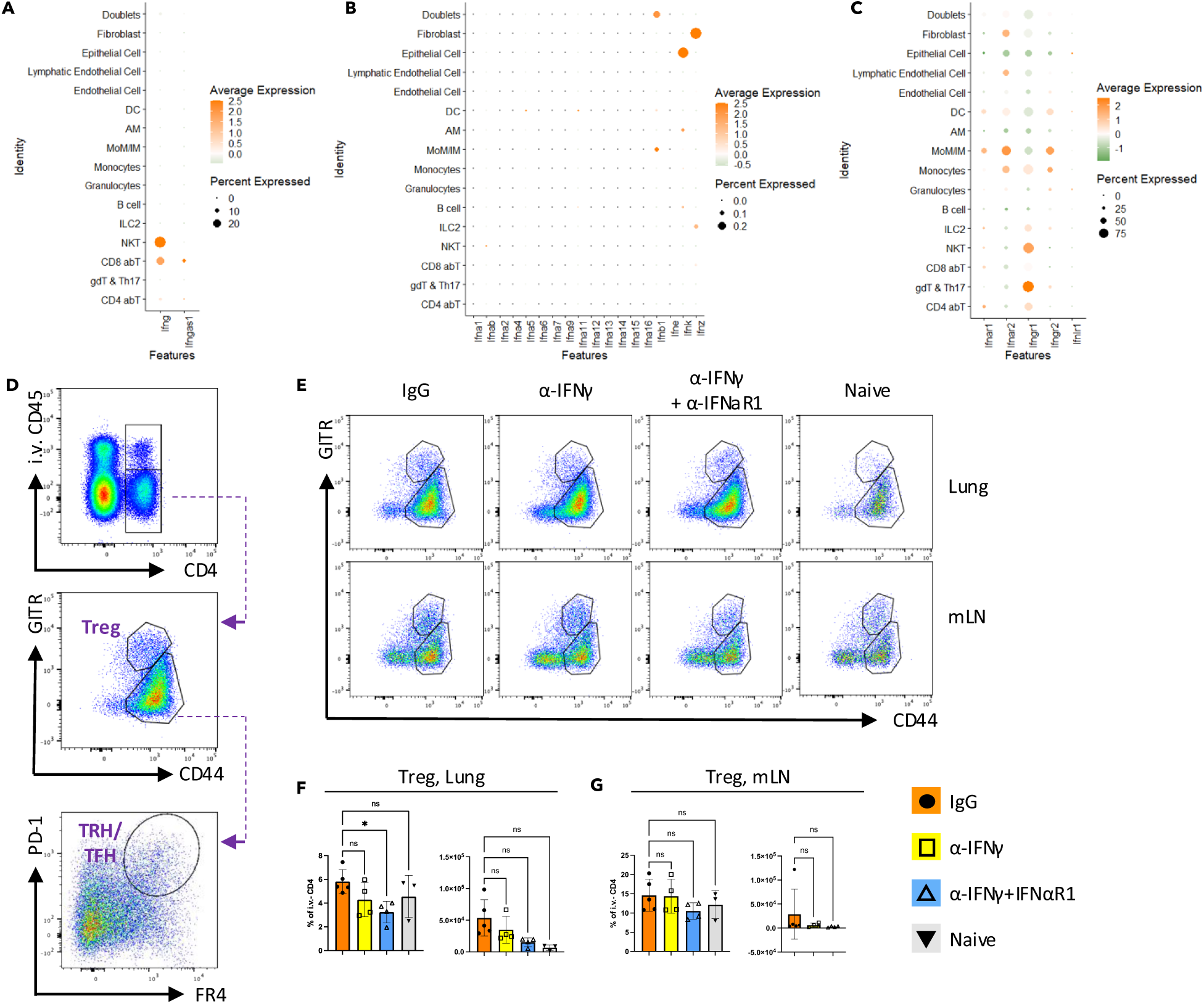
Exuberant type I and type II interferon signaling synergistically drives chronic sequelae in aged lungs. A. Dot plot displaying *Ifng* expression by defined cell types (Fig. 1C). Dot size and color represents the percentage of cells and average expression level, respectively, in given cell types (rows) and genes (columns). B. Dot plot illustrating expression of type I IFN genes in defined cell types (Fig. 1C). Dot size and color represents the percentage of cells and average expression level, respectively, in given cell types (rows) and genes (columns). C. Dot plot showing mRNA levels of type I and type II IFN receptors across defined cell types (Fig. 1C). Dot size and color represents the percentage of cells and average expression level, respectively, in given cell types (rows) and genes (columns). D. Gating strategy for Tregs and T_RH_/T_FH_ cells. E. Representative plots of Tregs in the lung and mLN for each experimental group. F. Bar graph quantifying Tregs in the lung. G. Bar graph quantifying Tregs in the mLN. In (F, G), each dot represents one animal. Statistical analysis was performed using repeated measures (RM) one-way ANOVA with Geisser-Greenhouse correction and multiple comparisons. Significance is indicated as *, p < 0.05; **, p < 0.01; ***, p < 0.001; ****, p < 0.0001.

**Supplementary Fig. 7.**
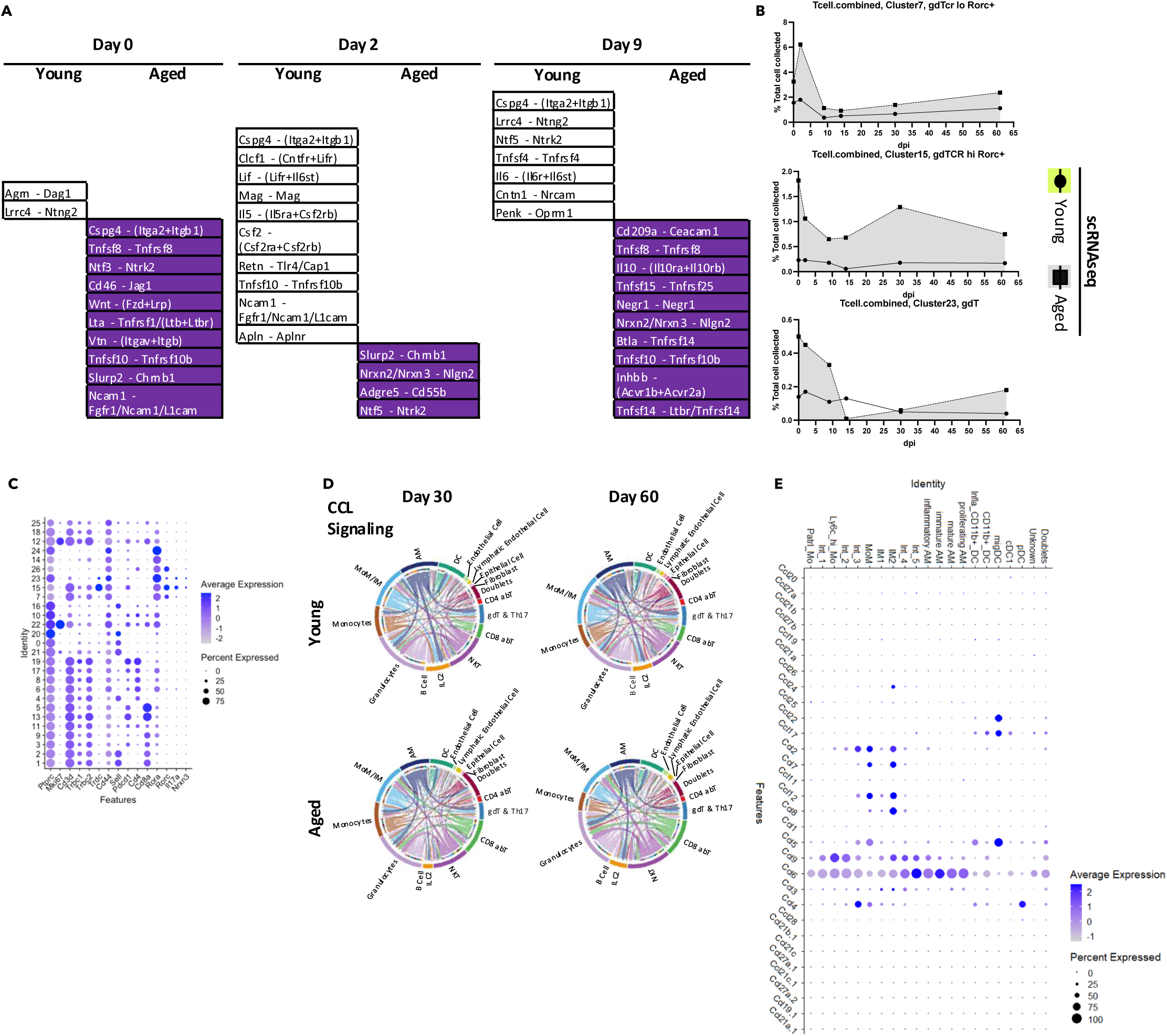
Cell-cell interaction analyses reveal unique pathways in young and aged hosts, and potential interactions between IMs and CD4^+^ T cells. A. Unique interactions identified in young or aged hosts at 0, 2, and 9 d.p.i. B. Kinetics of *Rorc* (RORγt) ^+^ cells quantified from scRNAseq data (Fig. 2A), expressed as a proportion of all events passing quality control in Fig. 1C. C. Dot plot showing selected features used to define subsets of *Rorc* (RORγt)^+^ cells. Dot size and color represents the percentage of cells and average expression level, respectively, in given clusters (rows) and genes (columns). D. Chord plot illustrating CCL signaling in young and aged hosts during the memory phase. Each arrow indicates an inferred ligand-receptor interaction. Each arrow originates from the signal-sending cell type and points toward the signal-receiving cell type. Arrow color represents the identity of the signal-sending cell, while arrow thickness reflects the inferred interaction strength. E. Dot plot displaying ligands involved in CCL signaling within MNPs (Fig. S4D). Dot size and color represents the percentage of cells and average expression level, respectively, in given genes (rows) and cell types (columns).

## Materials and Methods

### Mouse, infection, and antibody administration

Aged (∼24-month-old) wild-type (WT) female C57BL/6 mice were obtained from the National Institute on Aging or purchased from The Jackson Laboratory. Young (2-3-month-old) WT female C57BL/6 mice were also purchased from The Jackson Laboratory and subsequently bred in-house. Ccr2^icre^ ^Ai6^ mice were generated by breeding CCR2-CreER-GFP mice (RRID:IMSR_JAX:035229) with Ai6 mice (RRID:IMSR_JAX:007906), both obtained from The Jackson Laboratory. Mice of both genders were included in the transgenic experiments. To compare young and aged mice, cage bedding was exchanged once per week for four consecutive weeks prior to infection. All mice were housed in a specific pathogen-free environment. Animal experiments were approved by the Institutional Animal Care and Use Committees (IACUC) at the Mayo Clinic or the University of Virginia.

Mice were infected following a previously described protocol(Sun et al. 2009). After anesthesia, they were inoculated with mouse-adapted influenza A/PR/8/34 (∼75 PFU, a dose sublethal for young mice but lethal for approximately 50% of aged mice) in FBS-free DMEM (Corning).

Antibody administration to block IFNγ ± IFNαR in vivo was performed intraperitoneally (i.p.) once per week, starting at approximately 14 days post-infection (d.p.i.). Mice were treated with 200 μg/mouse anti-mouse IFNγ antibody (Clone XMG1.2, Bio X Cell) ± 250 μg/mouse anti-mouse IFNAR-1 antibody (Clone MAR1-5A3, Bio X Cell) or an InVivoMAb rat IgG1 isotype control (Clone HRPN, Bio X Cell).

### Tamoxifen treatment

To induce CRE expression in Ccr2^icre^ ^Ai6^ mice, tamoxifen was prepared with solvent consisting of sunflower oil and absolute ethanol (10:1 in volume). Mice received 1.5 mg/ms/day of Tamoxifen through intraperitoneal injection daily for 5 consecutive days starting 12-13 d.p.i. when their body weight started to recover.

### Broncho-alveolar lavage (BAL) fluid

Mice were humanely euthanized with an overdose of ketamine/xylazine. The skin was incised, and the thyroid gland along with surrounding connective tissues were carefully removed to expose the trachea. A small incision was then made in the trachea to insert a pipette tip configuration consisting of a 20 μl tip fitted into a 1000 μl tip. To obtain a relatively concentrated supernatant, the first lavage was performed with 600 μl of sterile PBS. After centrifugation at 1600 rpm for 5 minutes, the supernatant was aspirated and stored. The remaining cells were combined with the cells recovered from three subsequent lavages, each performed with 1000 μl of sterile PBS.

Red blood cells were lysed using ammonium-chloride-potassium (ACK) buffer (deionized water containing 0.15 M NH4Cl, 1 mM KHCO3, and 0.1 mM Na2EDTA). Following two washes with MACS buffer (PBS supplemented with 2% FBS and 2 mM/L Na2EDTA), the cell pellet was resuspended and prepared for flow cytometry.

### Tissue harvest and lung histopathology

To distinguish tissue resident and vasculature-associated immune population, 2μg/ms anti-mouse CD45 antibody (Biolegend, Clone: 30-F11) was injected intraperitoneally 5min before humanely euthanizing the animal. Lungs were perfused with 20 ml PBS through pulmonary circulation starting from right ventricle. A pair of hemostatic forceps was used to clamp the right main bronchi before harvesting the right lung in preparation of single-cell suspension. The left lung was subsequently inflated with 1ml 10% Paraformaldehyde for 30s before excision and immersed in 10% Paraformaldehyde for 48 hours. They were then transferred to ethanol (70%) and shipped to Mayo Clinic Histology Core Lab (Scottsdale, AZ) for histopathology. They were embedded in paraffin and 5 μm sections were cut for Hematoxylin and Eosin (H&E) or Masson’s trichrome staining.

For Lung single cell suspension preparation, lung tissue was mined into fine pieces and put in digestion buffer (IMDM with 183 U/ml type 2 collagenase (Worthington)). After incubation at 37 C° for 30min, the homogenate was put through the “m_lung_02” program on a gentleMACS tissue disrupter (Miltenyi). The suspension was then filtered through 70 μm mesh (Falcon). The flow through was palleted and washed with MACS buffer (PBS with 2% FBS and 2mM/L Na2EDTA), followed by lysis of red blood cells with ACK buffer. After 2 washes with MACS buffer, cells were ready for flow cytometry or scRNAseq library preparation.

### Flow cytometry

To perform surface staining, cells were first stained with viability dye (Zombie NIR™ Fixable Viability Kit, Cat# 423106; Zombie Aqua™ Fixable Viability Kit, Cat# 423102). Following the Fc receptor blockade with anti-mouse CD16/CD32 (Clone: 2.4G2, Bio X Cell), cells were incubated in MACS-buffer-based solutions with antibodies targeting surface markers on ice for 30min. If no intracellular proteins of interest exist in the panel, the samples would be ready after washing.

To stain transcription factors like FOXP3 or BCL6, Foxp3/Transcription Factor Staining Buffer Kit (Tonbo Biosciences) was utilized. Cells were incubated in the Fix/Perm buffer for 30min at room temperature in dark, and subsequently permeabilized in permeabilization buffer at 4 C° for 1h.

After staining in antibody containing permeabilization buffer for another 1h at 4 C°, cells were washed with PBS before ready for flow cytometry.

Data acquisition was enabled by 14-color Attune 742 NXT system (Life Technologies) or Cytek Aurora Borealis 5 laser Spectral Flow Cytometer. Data analysis was enabled by FlowJo (v10.10.).

### scRNAseq

To generate single cell RNA sequencing library, Chromium Next GEM Single Cell 5’ Library & Gel Bead Kit was utilized. Targeted cell number chosen was 10,000 cells/sample. CellRanger was utilized for annotation and alignment of the sequenced data, followed by integration and clustering with Seurat (v4). Cells with 200-6000 genes detected per cell together with less than 5% of mitochondria-related genes were included in further analysis. For better cell type identification, integration and clustering of the cells were then performed with graph-based anchoring (Stuart et al. 2019) between libraries from young and aged mice. In brief, differential expressed genes were identified separately. Intersection between the differential expressed genes was subsequently identified and utilized as “anchors” for integration. Unsupervised clustering was then performed (Fig1C). Myeloid cells and Lymphocytes were subsetted based on features in each cluster (FIgS1B). Integration and clustering were then repeated with the process described above.

Pathway analysis was enabled by clusterProfiler (Wu, Hu, et al. 2021) with Molecular Signatures Database (MSigDC) created by UC San Diego and Broad Institute (Subramanian et al. 2005; Mootha et al. 2003). Differential expressed genes were identified between datasets from young and aged mice in each time point, followed by rank-based enrichment score calculation. Gene list from selected pathways were then downloaded and used as input for scoring with AddModuleScore() function as a second layer of evaluation for relative expression level (Fig1D & 5A).

Cell Cycle Analysis was performed with built-in gene lists for S phase and G2/M phase in Seruat. Principal Component Analysis was performed using these two datasets and cells were assigned to “S phase” or “G2/M phase” based on “enrichment”, if cells were not considered “enriched” in any of these datasets, they were assigned to “G1/G0 phase”.

Bridging through Signac, trajectory Analysis of monocytes and macrophages were performed with Monocle3 (Trapnell et al. 2014). The cluster representing Ly6C^hi^ monocytes was chosen as “root”. Pseudotime values were then calculated and added to the Seurat object (Fig3B).

CellChat (Jin et al. 2021) was utilized to infer cell-cell interactions between all cells collected. Average gene expression of ligand/receptor from the built-in ligand-receptor interaction database (Cell-ChatDB, mouse) was calculated with Tukey’s trimean in each manually defined cell type (Fig1C). Among these ligand/receptors, 10% trimean was chosen as threshold for expression level to be considered as “over-expressed”. Probability of communication was quantified with the law of mass action. Significant interactions are determined through a statistical test involving the random permutation of cell group labels, followed by a recalculation of the probability of interaction.

## Mouse lung function measurement

Lung function measurements using FOT and the resulting parameters have been previously described (Narasimhan et al. 2024). In brief, animals were anesthetized with an overdose of ketamine/xylazine (100 and 10mg/kg intraperitoneally) and tracheostomized with a blunt 18-gauge canula (typical resistance of 0.18 cmH2O s/mL) secured in place with a nylon suture. Animals were then connected to the computer-controlled piston (SCIREQ flexiVent) and forced oscillation mechanics were performed under tidal breathing conditions with a positive-end expiratory pressure of 3 cm H2O. The measurements were repeated following thorough recruitment of closed airways (two maneuvers rapidly delivering TLC of air and sustaining the required pressure for several seconds, mimicking holding of a deep breath). Each animal’s basal conditions were normalized to their own maximal capacity. Measurement of these parameters before and after lung inflation allows for determination of large and small airway dysfunction under tidal (baseline) breathing conditions. Only measurements that satisfied the constant-phase model fits were used (>90% threshold determined by software). After this procedure, mice had a heart rate of ∼60 beats per minute, indicating that measurements were done on live individuals.

## Statistical analysis

The GraphPad Prism 9.0 (GraphPad Software) was executed for statistical analysis. The data is presented in the form of means ± SEM. The statistical methods for uncovering differences between groups were based on the Unpaired t-tests, one-way ANOVA, or two-way ANOVA according to the text. A p-value <0.05 was retained as a criterium for revealing statistical significance.

## Notes

### Competing Interest Statement

The authors have declared no competing interest.

